# InsP3R signaling mediates mitochondrial stress-induced longevity through actomyosin-dependent mitochondrial dynamics

**DOI:** 10.1101/2025.08.30.673265

**Authors:** Gaomin Feng, Elizabeth M. Ruark, Alexandra G. Mulligan, Eric K.F. Donahue, Anthony Hoang, Brianne Jacquet-Cribe, Li Peng, Kristopher Burkewitz

## Abstract

Certain forms of mitochondrial impairment confer longevity, while mitochondrial dysfunction arising from aging and disease-associated mutations triggers severe pathogenesis. The adaptive pathways that distinguish benefit from pathology remain unclear. Here we reveal that longevity induced by mitochondrial Complex I/*nuo-6* mutation in *C. elegans* is dependent on the endoplasmic reticulum (ER) Ca^2+^ channel, InsP3R. We find that the InsP3R promotes mitochondrial respiration, but the mitochondrial calcium uniporter is dispensable for both respiration and lifespan extension in Complex I mutants, suggesting InsP3R action is independent of matrix Ca^2+^ flux. Transcriptomic profiling and imaging reveal a previously unrecognized role for the InsP3R in regulating mitochondrial scaling, where InsP3R impairment results in maladaptive hyper-expansion of dysfunctional mitochondrial networks. We reveal a conserved InsP3R signaling axis through which calmodulin and actomyosin remodeling machineries, including Arp2/3, formin FHOD-1, and MLCK, constrain mitochondrial expansion and promote longevity. Disruption of actin remodeling or autophagy mimics InsP3R loss. Conversely, driving fragmentation ameliorates mitochondrial expansion and rescues longevity, supporting a model in which InsP3R-dependent actin remodeling sustains mitochondrial turnover. These findings establish an inter-organelle signaling axis by which ER calcium release orchestrates mitochondrial-based longevity through cytoskeletal effectors.

## Introduction

A variety of genetic and pharmacological approaches that perturb mitochondrial electron transport chain (ETC) function extend lifespan in worms, flies, rodents and potentially humans^1–7^. On the other hand, severe mitochondrial impairment results in toxicity and mitochondrial disease. The cellular adaptations that determine whether mitochondrial perturbation results in enhanced resilience versus pathogenesis thus possess enormous therapeutic potential, but remain surprisingly unclear. Previously, the search for mechanisms of mitochondrial stress-induced longevity has yielded insights on roles for signaling from reactive oxygen species (ROS), metabolic rewiring, and retrograde signals to the nucleus that remodel chromatin and transcriptional outputs, such as the mitochondrial unfolded protein response (UPRmt)^8–10^. However, outputs such as the UPRmt and ROS levels do not consistently correlate with lifespan and are not sufficient for longevity, indicating that mechanistically important adaptive mechanisms remain to be discovered^9,11–13^. Adding further complexity, perturbing different components of the ETC can promote longevity through distinct mechanisms^14,15^. Even within the same complex, different types of perturbation—such as mutation versus knockdown—may engage genetically separable pathways that promote longevity, the basis for which remains poorly understood^16^. Thus, identifying pathways specifically required for longevity in defined mitochondrial stress contexts is critical.

The current challenges in explaining how mitochondrial stress leads to life-extension has led to suggestions that adaptive remodeling of processes outside of mitochondria must be playing important, poorly understood roles in mediating longevity^9^. Along these lines, one underexplored avenue in understanding mitochondrial longevity paradigms relates to central mitochondrial roles in buffering and responding to fluctuations in intracellular Ca^2+^ levels. Mitochondrial impairment results in altered calcium uptake and storage in the matrix^17–20^, thus leading to potential broadscale rewiring of calcium signaling processes in other cellular compartments^21^. Indeed, dysregulation of cellular calcium handling and mitochondrial dysfunction commonly co-occur in age-related diseases and are thought to antagonize one another^22,23^, pointing to calcium-dependent processes as key determinants of healthy vs. pathology-associated aging trajectories. How aberrant intracellular calcium signaling exacerbates mitochondrial dysfunction in these contexts remains unclear, however. This lack of clarity stems in part from the inherent challenges in dissecting the multifaceted and highly interconnected interplay between intracellular Ca^2+^ signaling and mitochondrial homeostasis, which involves links to bioenergetic control in the matrix and multiple pathways regulating mitochondrial dynamics and organization.

Inositol triphosphate receptor (InsP3R) channels on the endoplasmic reticulum (ER) membrane act as a unique control point for intracellular calcium signaling, as they are ubiquitously expressed in metazoans and linked directly to both mitochondrial and cytosolic Ca^2+^ pools. InsP3R channels are enriched at ER-mitochondrial contact sites and capable of controlling bioenergetic tone through direct Ca^2+^ flux into the mitochondrial matrix^20,24–26^. The mitochondrial outer membrane is relatively permeant to Ca^2+^ due to high levels of voltage-dependent anion channels (VDAC), while the mitochondrial calcium uniporter (MCU) mediates low-affinity, high-capacity uptake of Ca^2+^ into the matrix. Notably, MCU Ca^2+^ uptake is sensitive to mitochondrial membrane potential, and mitochondrial stress can result in cytosolic Ca^2+^ spillover that can reciprocally modulate activity of the InsP3R^17–19,21^, thus speculatively providing an avenue for InsP3R channels to sense and respond to mitochondrial dysfunction. In vertebrates, InsP3R-dependent increases in matrix Ca^2+^ stimulate the activity of multiple dehydrogenases associated with the tricarboxylic acid (TCA) cycle, including pyruvate dehydrogenase (PDH), isocitrate dehydrogenase, oxoglutarate dehydrogenase, and potentially even the ETC itself^27^.

Beyond bioenergetic stimulation, regulation of mitochondrial dynamics has emerged as another major factor in determining the outcome of mitochondrial stress responses and aging outcomes^28^. Recent studies have revealed that calcium plays important roles in activating mitochondrial fission machineries^29,30^ and actin remodeling processes in response to mitochondrial damage^31–33^. Notably, these mitochondrial actin networks cage and segregate depolarized mitochondria from the larger network, alter regulation of mitochondrial motility and fission/fusion dynamics, and help to promote adaptive metabolic shifts, such as glycolytic upregulation^31–34^. Generally, these studies have focused primarily on acute damage paradigms *in vitro*, however, and little is known about whether calcium-dependent remodeling of mitochondrial networks plays causal roles in organismal aging or mitochondrial longevity paradigms.

Here we employed *C. elegans* to test whether inter-organelle Ca^2+^ signaling plays a role in the adaptive benefits of mitochondrial stress. We focus on the InsP3R, a conserved ER Ca²⁺ channel linked to mitochondrial homeostasis, but whose physiological role in aging and stress adaptation is unclear. We demonstrate that InsP3R activity is required for the longevity conferred by Complex I dysfunction, and that this effect occurs independently of matrix Ca²⁺ uptake through the MCU. Instead, our transcriptomic, imaging, and genetic analyses reveal that InsP3R signaling coordinates calcium-sensitive actomyosin remodeling, which constrains maladaptive mitochondrial expansion and enables quality control to generate a permissive landscape for successful adaptation to mitochondrial dysfunction. These findings identify a previously unrecognized role for cytosolic Ca²⁺ signaling and actin dynamics in mitochondrial longevity and provide a framework for understanding how mitochondrial dysfunction and calcium dysregulation interact during aging.

## Results

### InsP3R signaling is essential for adaptation and lifespan extension following mitochondrial perturbation

The roles of calcium signaling in mitochondrial longevity remain poorly defined. Here we set out to test the hypothesis that ER-mitochondrial calcium crosstalk may act as an important driver of longevity in these contexts. To modulate this crosstalk in the aging model, *C. elegans*, we focused on the InsP3R, which is closely linked to both mitochondrial bioenergetics in vertebrates and to longevity in *C. elegans* in the contexts of epidermal growth factor (EGF) signaling and ER stress^35,36^. While mammals possess three InsP3R genes, *C. elegans* possess only one, *itr-1*, presenting a simplified system for genetic modulation. While complete loss of InsP3R/*itr-1* function compromises viability^37^, a temperature-sensitive point mutant, *itr-1(sa73)* exhibits mild to moderate reductions in the frequency, velocity and magnitude of physiologic Ca^2+^ oscillations under normal culture conditions (20 °C)^38,39^. Consistent with previous links to bioenergetic signaling in worms and direct bioenergetic regulation in vertebrates^25,36^, we found that InsP3R/*itr-1(sa73)* mutants exhibit reductions in basal and FCCP-induced oxygen consumption rates (OCR) that are similar in magnitude to animals harboring a mutation in Complex I subunit, *gas-1* (Fig. 1A-C). This effect on OCR is independent of gross ER stress or dysfunction, as impairing ER homeostasis more directly through the *ire-1*/*xbp-1* pathway has no effect on respiration (Extended Data Fig. 1A-C). These results indicate a key role for InsP3R/*itr-1* signaling in maintaining mitochondrial respiration at the organismal scale.

**Figure 1:**
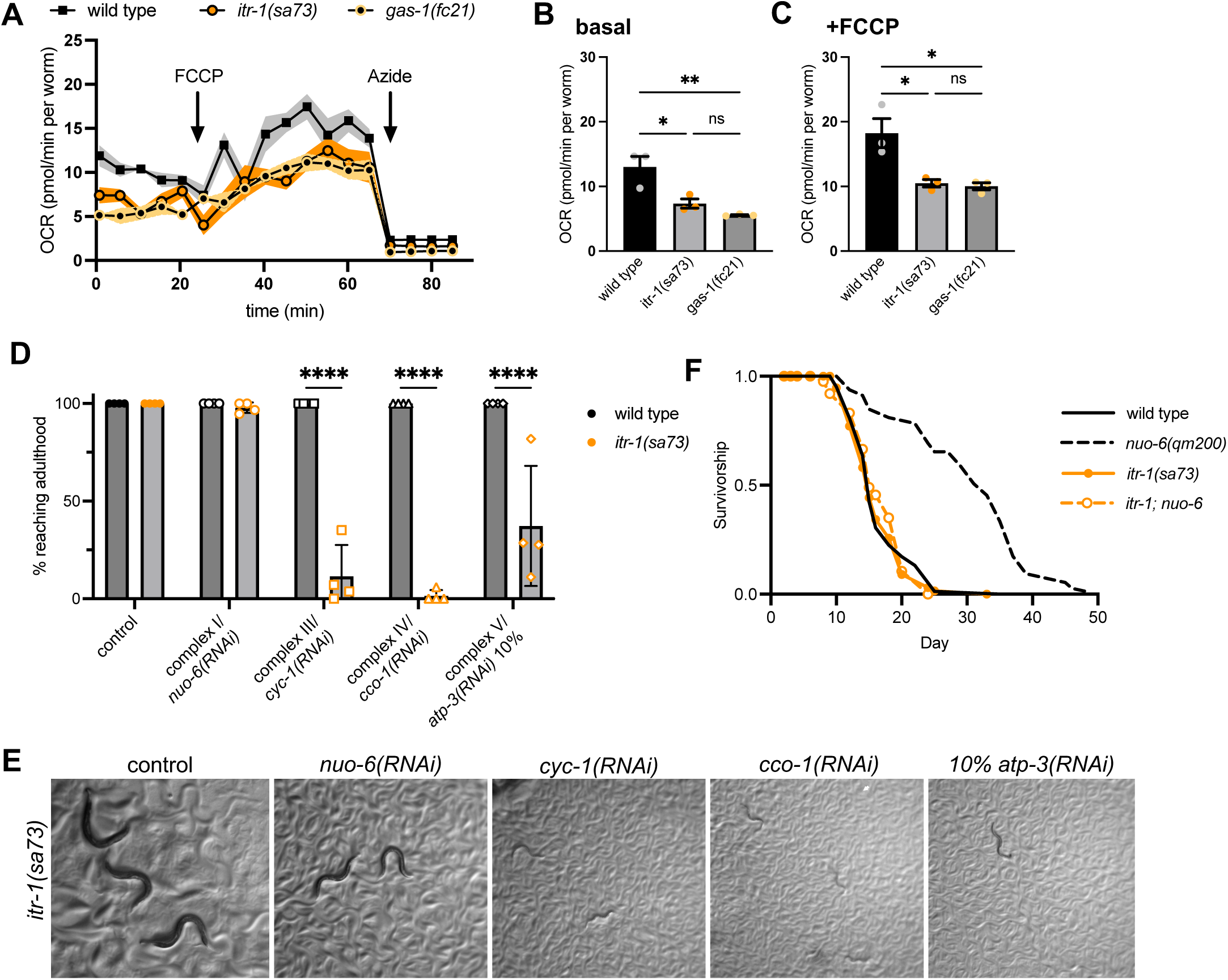
InsP3R/*itr-1* is required for longevity in Complex I/*nuo-6* mutants. (A) Representative traces of OCR in InsP3R/*itr-1* and *gas-1* mutants; mean ± SEM. Additions of FCCP and sodium azide are indicated by arrows. (B, C) Quantification of basal (B) and FCCP-induced (C) OCR; n = 3 independent trials; mean ± SEM; one-way ANOVA with Tukey’s multiple comparisons test. (D) Percent of wild type and InsP3R/*itr-1* mutant worms reaching adulthood 10 days post egg lay after dsRNA knockdown against long-lived ETC complexes. n = 110-211 worms per condition scored across 4 independent trials; mean ± SD; two-way ANOVA with Fisher’s LSD test. (E) Representative images showing the development of *itr-1(sa73)* worms 10 days post egg lay on RNAi against ETC complexes. All images taken at 4x. (F) Lifespan of InsP3R/*itr-1* and Complex I/*nuo-6* double mutants. n = 100 worms per condition. Statistical significance of lifespan curves was determined by a log-rank Mantel-Cox test. See Supplementary Table 1 for lifespan statistics. ns p > 0.05; * p < 0.05; ** p < 0.01; and **** p < 0.0001.

To investigate if InsP3R/*itr-1* signaling is broadly required for organismal adaptation to mitochondrial dysfunction, we first tested how InsP3R/*itr-1* mutants tolerate RNAi-mediated impairment of ETC complexes I (*nuo-6)*, III (*cyc-1*), IV (*cco-1*), and V (*atp-3*). Intriguingly, we found that knockdown of Complexes III, IV, and V caused profound developmental arrest selectively in the InsP3R/*itr-1* background (Fig. 1D,E). In contrast, Complex I/*nuo-6* knockdown was compatible with development in these animals. These differences were not attributable to RNAi efficacy (Extended Data Fig. 1D,E), and instead suggest that InsP3R function is essential for developmental adaptation to ETC inhibition at Complexes III–V. Because Complex I/*nuo-6* impairment was uniquely tolerated in InsP3R/*itr-1* mutants, we next asked whether InsP3R function was also required for the lifespan extension typically observed in Complex I mutants. We crossed *itr-1(sa73)* into a long-lived *nuo-6(qm200)* background^16^ and found that reduced InsP3R function completely suppressed the extended lifespan of *nuo-6* mutants (Fig. 1F), which typically ranges from 75–120% on standard media (see Supplementary Table 1 for complete statistics). This genetic suppression supports a model in which InsP3R signaling is required for the adaptive responses that mediate longevity in the context of Complex I impairment.

To determine whether this requirement was specific to the *nuo-6* mutant or extended to other Complex I–related longevity paradigms, we also tested the effect of InsP3R loss on lifespan extension induced by RNAi knockdown of *nuo-6*. Prior work demonstrated that *nuo-6(RNAi)* and *nuo-6* mutation extend lifespan via genetically independent mechanisms, though those precise mechanisms remain undefined^16^. In line with this, we found *nuo-6(RNAi)* longevity was unaffected by InsP3R/*itr-1* (Extended Data Fig. 1F), indicating that InsP3R signaling is not globally required for ETC-induced longevity. To further assess the specificity of InsP3R’s requirement for longevity, we tested two other paradigms distinct from ETC perturbation. First, lifespan extension from *age-1* insulin-signaling mutants was only modestly suppressed by InsP3R/*itr-1* loss (Extended Data Fig. 1G), and dietary restriction robustly extended lifespan in both wild type and InsP3R/*itr-1(sa73)* mutants (Extended Data Fig. 1H). These findings demonstrate that InsP3R/*itr-1* mutants are capable of mounting adaptive longevity responses in several contexts, and suggest that InsP3R signaling is selectively required for the pro-longevity program triggered by Complex I mutation.

Finally, we also used an independent approach for reducing InsP3R function without potential disruption of channel structure in InsP3R point mutants. We generated animals ubiquitously expressing an InsP3 sponge, a validated construct that sequesters InsP3R agonist to reduce InsP3R-driven calcium efflux^37^. In this model, we observed that lifespan extension was substantially diminished in the sponge background (∼75-115% in controls ∼40% when the sponge is expressed (Extended Data Fig. 1I), supporting a causal role for InsP3R signaling per se in mitochondrial stress-induced longevity.

### InsP3R-dependent control of respiration and lifespan is independent of MCU and matrix metabolic processes

We next sought to understand the mechanism by which InsP3R function promotes longevity in the context of Complex I inhibition. In cancer cells, constitutive InsP3R-dependent stimulation of mitochondrial bioenergetics sustains the metabolic demands of rapid proliferation, and impaired ER-mitochondrial Ca^2+^ flux results in “bioenergetic catastrophe”^42^ that compromises cell viability. A similar threshold-dependent phenomenon exists in *C. elegans*, where milder impairment of ETC function promotes lifespan extension, but stronger inhibition triggers dysfunction and early death^25,40^. Drawing on this parallel, we hypothesized that InsP3R-mediated Ca^2+^ transfer to stressed mitochondria could sustain mitochondrial metabolic functions above a critical threshold, thereby enabling adaptive responses and supporting longevity. Consistent with a model of bioenergetic catastrophe, we found that combining Complex I/*nuo-6* and InsP3R/*itr-1* mutations reduced basal and FCCP-induced OCR to lower levels than either of the single mutants (Fig. 2A-C). This additive impairment of respiration in Complex I; InsP3R double-mutants supports the concept that InsP3R signaling either enhances or acts in parallel of Complex I to promote mitochondrial functions.

**Figure 2:**
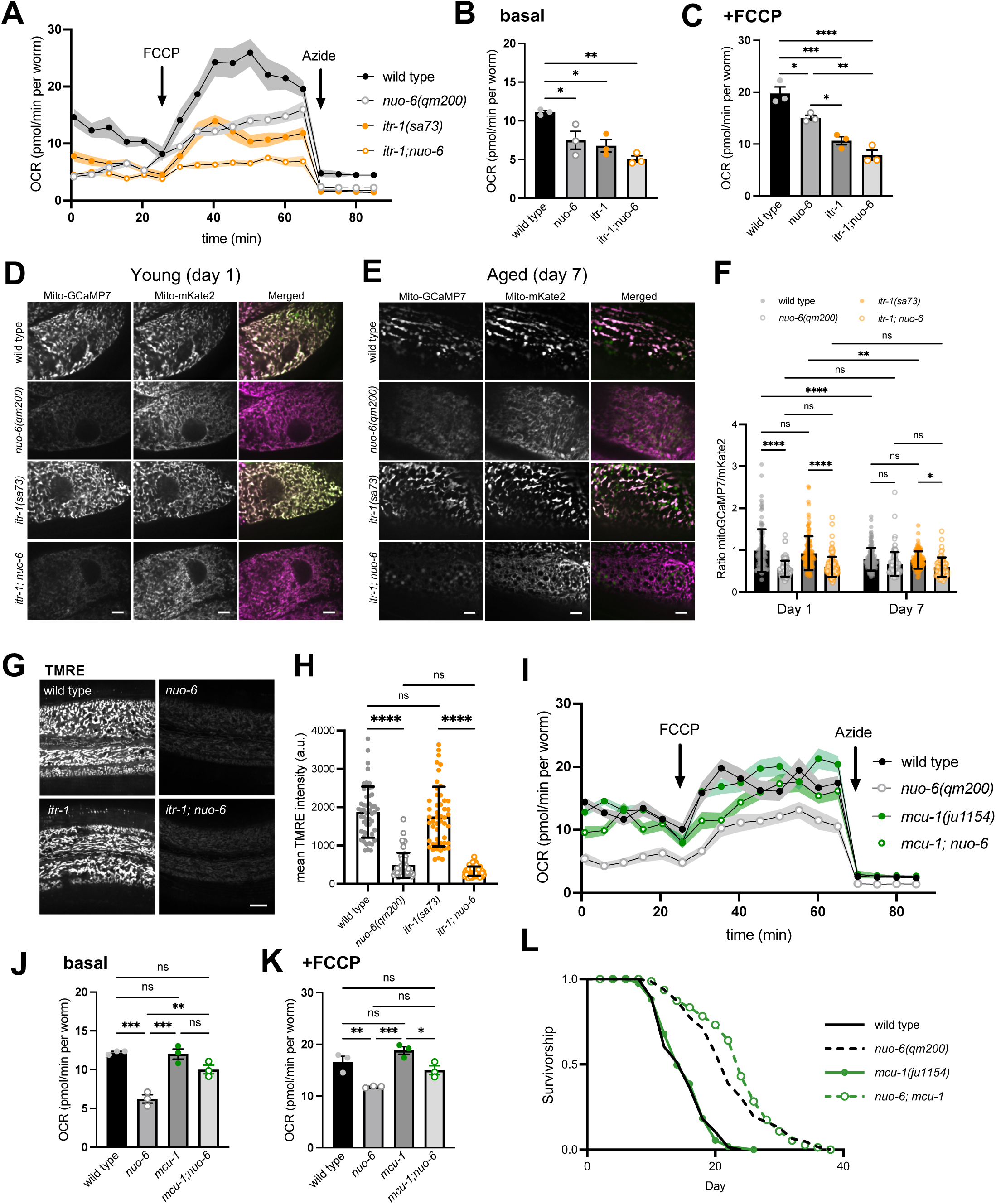
InsP3R roles in Complex I/*nuo-6* longevity are uncoupled from MCU and matrix calcium. (A) Representative OCR traces in InsP3R/*itr-1* and Complex I/*nuo-6* mutants; mean ± SEM. Additions of FCCP and sodium azide are indicated by arrows. (B, C) Quantification of basal (B) and FCCP-induced (C) OCR in InsP3R/*itr-1* and Complex I/*nuo-6* mutants. n = 3 independent trials; mean ± SEM; one-way ANOVA with Tukey’s multiple comparisons test. (D, E) Representative images of Mito-GCaMP7 and mito-mKate2 imaging in intestine of day 1 (D) and day 7 (E) adults of InsP3R/*itr-1* and Complex I/*nuo-6* mutants. Scale bar: 5 µm. (F) Quantification of the ratio of GCaMP7 to mKate2 fluorescence as an indicator of basal mitochondrial calcium level. n = 115, 115, 123, 122, 138, 103, 120, 59 worms from 3 independent trials (left to right); mean ± SD; two-way ANOVA with Tukey’s multiple comparisons test. (G) Representative images of TMRE staining in hypodermal mitochondria of day 1 adults of InsP3R/*itr-1* and Complex I/*nuo-6* mutants. Scale bar: 10 µm. (H) Quantification of mean TMRE intensity. n = 50, 50, 50, 59 worms from 3 independent trials (left to right); mean ± SD; one-way ANOVA with Tukey’s multiple comparisons test. (I) Representative OCR traces in Complex I/*nuo-6* and *mcu-1* mutants; mean ± SEM. Additions of FCCP and sodium azide are indicated by arrows. (J, K) Quantification of basal (J) and FCCP-induced (K) OCR in Complex I/*nuo-6* and *mcu-1* mutants. n = 3 independent trials; mean ± SEM; one-way ANOVA with Tukey’s multiple comparisons test. (L) Lifespan of Complex I/*nuo-6* and *mcu-1* mutants. n = 100 worms per condition. Statistical significance of lifespan curves was determined by a log-rank Mantel-Cox test. See Supplementary Table 1 for lifespan statistics. ns p > 0.05; * p < 0.05; ** p < 0.01; *** p < 0.001; and **** p < 0.0001.

Based on previous observations in cancer cells^30,43,44^, we next tested the hypothesis that InsP3R signaling potentiates Complex I/*nuo-6* dependent longevity and mitochondrial bioenergetics via MCU-mediated Ca^2+^ flux and downstream processes in the matrix. First, we asked whether matrix [Ca^2+^] correlate with the effects of Complex I/*nuo-6* and InsP3R/*itr-1* on lifespan. To quantify relative matrix Ca^2+^ levels, we co-targeted Ca^2+^-sensitive GCaMP7 and Ca^2+^- insensitive mKate2 as a normalization factor to the mitochondrial matrix (Fig. 2D-F) after confirming the reporter responded as expected to conditions that reduce (*mcu-1* loss) or enhance (SERCA/*sca-1* inhibition) matrix Ca^2+^ levels (Extended Data Fig. 2A,B). We observed robust decline of resting matrix Ca^2+^ in Complex I*/nuo-6* mutants and no additive effects when InsP3R and Complex I mutants were combined (Fig. 2D-F). As mitochondrial Ca^2+^ uptake is membrane potential-dependent, we tested the effects of Complex I/*nuo-6* and InsP3R/*itr-1* mutants on mitochondrial membrane potential via tetramethylrhodamine ethyl ester (TMRE) staining. Consistent with the observed changes in matrix Ca^2+^, we observed a substantial decrease in TMRE staining of mitochondria in Complex I/*nuo-6* mutants and no apparent contributions from InsP3R/*itr-1* (Fig. 2G,H). These results highlight the substantial alteration in intracellular calcium handling in mitochondrial longevity mutants, while suggesting the capacity for InsP3R-to-matrix Ca^2+^ flux is limited by reduced membrane potential in these contexts.

While resting matrix [Ca^2+^] does not correlate with OCR or the lifespan outcomes, we also reasoned that difficult-to-detect Ca^2+^ transients from the InsP3R could still stimulate bioenergetics and other mitochondrial behaviors^30,43,44^. We first examined whether calcium-sensitive matrix dehydrogenases are responsive to InsP3R signaling in *C. elegans.* In vertebrates, rises in matrix Ca^2+^ levels stimulate PDH phosphatase activity, subsequently leading to dephosphorylation of PDH subunit E1α at S293^45^, a site fully conserved in *C. elegans pdha-1*. While Western blots of phospho-PDH revealed higher levels of the inactive, phosphorylated PDH enzyme in InsP3R/*itr-1* mutants (Extended Data Fig. 2C,D), potentially consistent with reduced phosphatase activity, we also observed an increase in total PDH levels (Extended Data Fig. 2E,F), suggesting that an expansion of mitochondrial enzymes or networks in InsP3R/*itr-1* mutants may drive the effect on phospho-PDH levels rather than a strong shift towards inactivation of PDH. Finally, to rule out potential roles for matrix transients and the MCU complex more generally, we incorporated deletion mutants of *mcu-1*, the sole channel-forming MCU subunit in *C. elegans*. While *mcu-1* mutants exhibited reduced resting [Ca^2+^]_m_ as expected (Extended Data Fig. 2A), but surprisingly we found that animals lacking MCU have no observable reduction in OCR (Fig. 2I-K). Furthermore, lifespan extension of Complex I mutants is unaffected by MCU functionality (Fig. 2L). Taken together, these results indicate that while the InsP3R stimulates mitochondrial respiration at the organismal scale in *C. elegans*, the mechanism by which InsP3R signaling supports mitochondrial adaptation and Complex I longevity is largely independent of direct transfer of Ca^2+^ from ER stores to the matrix via MCU.

### Transcriptomics reveal dysregulation of mitochondrial biogenesis in InsP3R mutants under mitochondrial stress

We next took an unbiased approach to identify candidate mechanisms by which the InsP3R supports longevity in Complex I mutants. We performed bulk RNA-Seq analysis of a panel of mutants that included wild type controls, InsP3R/*itr-1*, Complex I/*nuo-6,* and InsP3R/*itr-1*; Complex I/*nuo-6* double-mutants. We defined differentially expressed (DE) transcripts as those which exhibited a fold-change (FC) of at least 1.5x (P_adj_ < 0.01) relative to wild type. This approach revealed widespread transcriptional changes, yielding >2,000 DE transcripts in each mutant (Fig. 3A). We categorized DE transcripts to highlight the genes and cellular processes predicted to drive the lifespan phenotypes downstream of Complex I and InsP3R signaling (Fig. 3B). We reasoned the most promising candidates would fall into Category I: transcripts altered in Complex I mutants that are InsP3R/*itr-1* dependent. Additionally, Category II transcripts are those altered in Complex I mutants independently of InsP3R function, thus likely uncoupled from lifespan, while Category III transcripts exhibit a synthetic expression profile, altered only when both Complex I and InsP3R function are impaired (Fig. 3A,B & Supplementary Table 2).

**Figure 3:**
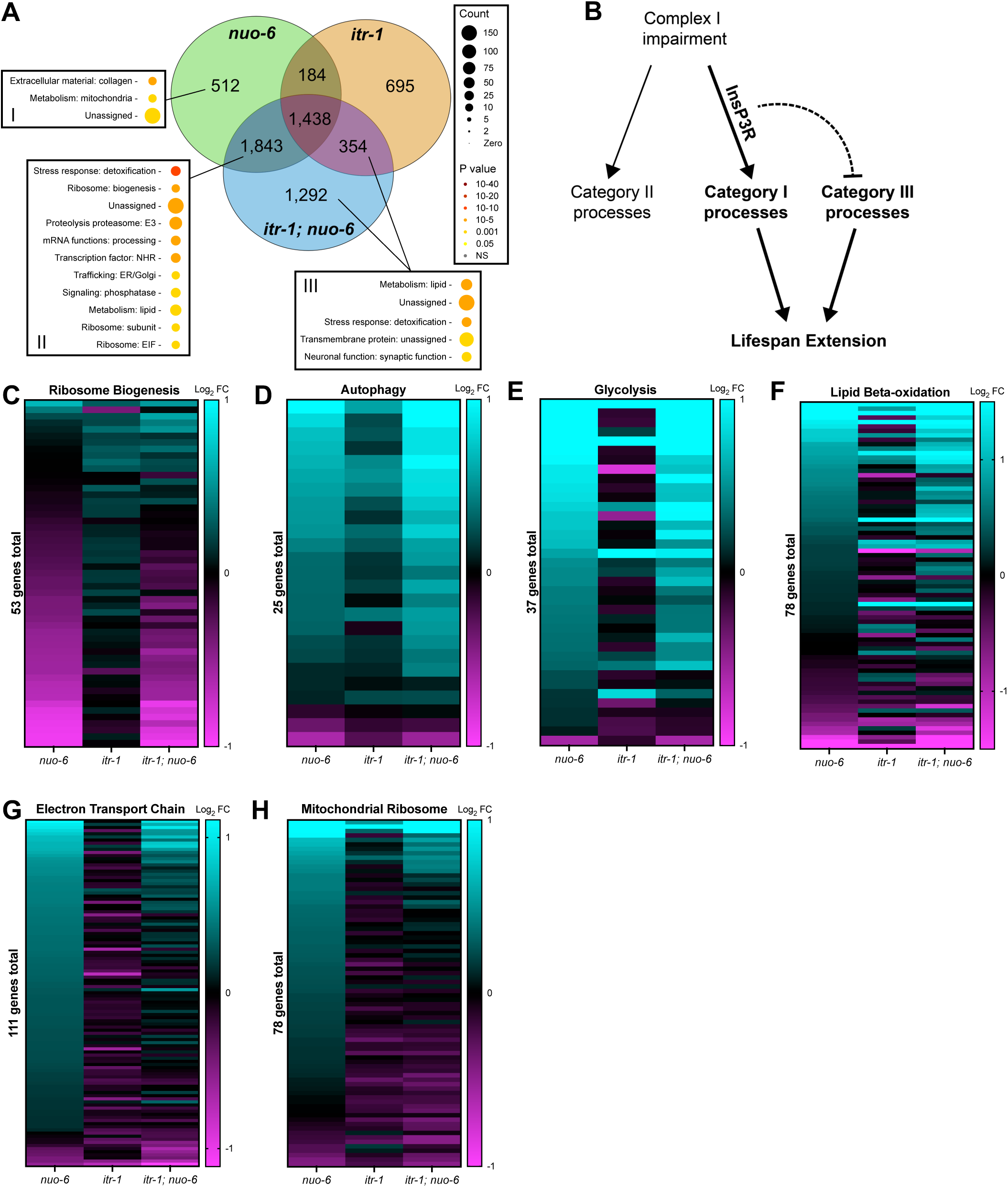
Transcriptomic analysis reveals InsP3R-dependent regulation of mitochondrial biogenesis and adaptive responses to Complex I impairment. (A) Venn diagram representing differentially expressed genes (Log_2_ FC of 1.5x, P_adj_ < 0.01) in Complex I/*nuo-6* (green), InsP3R/*itr-1* (orange), and InsP3R/*itr-1;* Complex I/*nuo-6* (blue). Boxes represent categories (WormCat – Category 2) of genes enriched in the indicated groups. See Supplementary Table 2 for summary of differentially expressed genes. (B) Model depicting how DE gene categories correlate with genetic regulation of lifespan by *nuo-6* and *itr-1*. (C-H) Heatmaps comparing the expression of mRNA transcripts in Complex I/*nuo-6*, InsP3R/*itr-1*, and InsP3R/*itr-1;* Complex I/*nuo-6* mutants for (C) ribosome biogenesis, (D) autophagy, (E) glycolysis, (F) lipid beta-oxidation, (G) electron transport chain, and (H) mitochondrial ribosome genes. All genes normalized to wild type.

Given the large-scale remodeling of gene expression we observed in these contexts, we next performed gene enrichment analysis within these categories^46–48^. Notably, our analysis of Complex I/*nuo-6* mutants closely matched established signatures of the UPRmt observed upon a variety of forms of mitochondrial stress^49,50^. Some of the strongest transcriptional changes we observed in Category II (Complex I dependent, InsP3R-independent) involved transcriptional modules associated with the UPRmt (Fig. 3A), including reduced translation and cytosolic ribosome biogenesis (Fig. 3C), upregulated protein turnover and autophagy (Fig. 3D), and upregulation of glycolysis (Fig. 3E). Interestingly, altered lipid metabolism was a signature present in both Category II and III, indicating that some aspects of lipid metabolism are remodeled similarly in Complex I mutants regardless of InsP3R status, while others are uniquely altered in InsP3R/*itr-1;* Complex I double-mutants (Fig. 3A). These results suggest that Complex I mutants and InsP3R signaling may converge on distinct sets of lipid metabolism genes. Consistent with this notion, we observed similar overall trends in lipid beta-oxidation transcripts between Complex I/*nuo-6* and InsP3R/*itr-1;* Complex I/*nuo-6* conditions, while a subset were strongly InsP3R-dependent (Fig. 3F).

Finally, the processes within Category I include those that are altered during Complex I dysfunction and dependent on intact InsP3R function (Fig. 3A,B), and which we therefore expected to be enriched for roles in mediating InsP3R-dependent lifespan extension. Strikingly, only 2 categories were enriched in this subset of transcripts: collagens and mitochondrial functions, strongly suggesting that InsP3R signaling was likely to be mediating adaptation and longevity in Complex I mutants at least in part through a mechanism involving mitochondrial homeostasis. Upon closer examination of the genes and processes within this Mitochondrial category, we observed robust induction of mitochondrial ETC components and mitochondrial ribosomes in Complex I/*nuo-6* mutants, indicative of a compensatory mitochondrial biogenesis response during chronic mitochondrial dysfunction. However, this induction was substantially blunted in InsP3R/*itr-1* mutants (Fig. 3G,H), revealing a novel role for InsP3R signaling in regulating adaptive expansion of the mitochondrial compartment.

Altogether, our analysis confirms overlap with prior mitochondrial stress signatures and indicates that certain UPRmt transcriptional modules—particularly those related to mitochondrial biogenesis and scaling—are selectively dependent on InsP3R function. Interestingly, InsP3R signaling emerged as a hit from two prior genetic screens for UPRmt modulators, though neither study pursued mechanistic validation^49,50^. Our data build on these findings by demonstrating that InsP3R function shapes specific transcriptional modules of the UPRmt during Complex I stress.

### InsP3R signaling controls mitochondrial network scaling in response to Complex I dysfunction

To explain the effects of InsP3R signaling on mitochondrial biogenesis-associated transcripts, we hypothesized that the InsP3R may play a role in mitochondrial network expansion and quality control during stress. We analyzed the impacts of InsP3R and Complex I/*nuo-6* mutants on mitochondrial networks directly via confocal microscopy of animals harboring natively labeled Complex IV subunit, COX-4::eGFP, focusing on the primary metabolic tissues of *C. elegans*, the hypodermis (Fig. 4) and intestine (Extended Data Fig. 3). Mirroring the transcriptional signature of upregulated biogenesis, the mitochondrial networks of Complex I/*nuo-6* mutants undergo robust, compensatory expansion and adopt a predominantly tubular network morphology, while InsP3R/*itr-1* mutants alone exhibit mild mitochondrial phenotypes (Fig. 4A-C). In a surprising contrast with the prediction from both our transcriptomic results and respiration measurements, however, we observed an exaggerated expansion of the mitochondrial network in InsP3R/*itr-1*; Complex I/*nuo-6* double-mutants (Fig. 4A-C). This effect was generally consistent in the intestine (Extended Data Fig. 3), where double-mutants display an expanded and highly fused mitochondrial footprint similar to Complex I mutants, but with further elevated levels of COX-4::GFP. As mitochondrial dynamics and function are highly dynamic during aging^50^, we followed this phenotype as the animals reached ages where subsets of mitochondria in control animals begin exhibiting characteristic fragmentation and swelling (Fig. 4D,E). Consistent with this role for the InsP3R in restraining mitochondrial expansion, Complex I/*nuo-6* driven mitochondrial expansion is enhanced over age in InsP3R/*itr-1* mutants (Fig. 4D-F). Indeed, the mitochondrial network phenotype in InsP3R/*itr-1*; Complex I/*nuo-6* double mutants becomes relatively extreme, adopting such a dense and hyperfused configuration that COX-4::eGFP mitochondria appear to fill the cytosol, punctuated only by nuclei and what appear to be accumulating lipid-containing vesicles (Fig. 4D). To better resolve these mitochondrial and vesicular structures, we performed transmission electron microscopy (TEM) imaging of transverse cross-sections (Fig. 4G-J). Consistent with the expanded tubular network observed in confocal images, we found mitochondrial cross-sections in the Complex I/*nuo-6* mutants that appear relatively densely packed together (Fig. 4G,I). In Complex I; InsP3R double-mutants, the density of individual mitochondrial tubules appears consistent, but the mitochondria are substantially larger in both young and aged animals, suggesting InsP3R mutants enhance or are additive to processes resulting in mitochondrial expansion (Fig. 4G-J). Additionally, TEM images indicated that the vesicles punctuating the hyper-expanded mitochondrial network in InsP3R mutants are consistent with lipid droplets (Fig. 4G,I), suggesting that defects in InsP3R signaling may result in lipid accumulation that increases with age.

**Figure 4:**
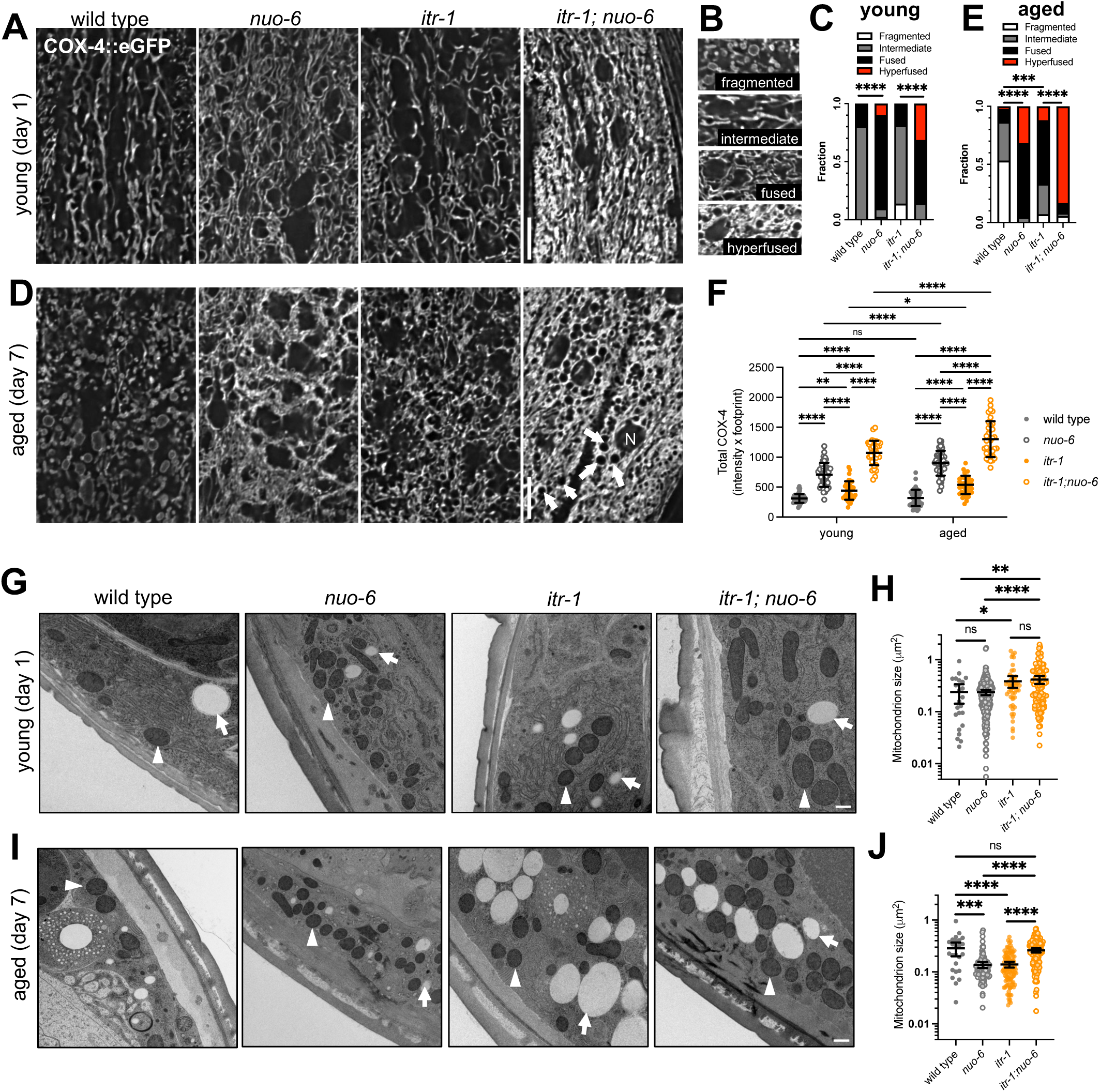
InsP3R signaling constrains mitochondrial network expansion during Complex I dysfunction. (A) Representative images of mitochondrial networks (COX-4::eGFP) in hypodermis of young (day 1) worms. Scale bar: 10 µm. (B) Representative examples of fragmented, intermediate, fused, and hyperfused mitochondrial morphologies. (C) Categorization of mitochondrial morphologies in young (day 1) worms. n = 46, 41, 43, 35 worms from 2 independent trials (left to right); Kruskal-Wallis test with Dunn’s multiple comparisons test. (D) Representative images of mitochondrial networks (COX-4::eGFP) in hypodermal cells of aged (day 7) worms. Scale bar: 10 µm. “N” = nucleus, arrows indicate lipid droplets. (E) Categorization of mitochondrial morphologies in aged (day 7) worms. n = 45, 44, 42, 36 from 2 independent trials (left to right); Kruskal-Wallis test with Dunn’s multiple comparisons test. (F) Quantification of total COX-4 (mean COX-4::eGFP intensity x mitochondrial footprint) in young (day 1) and aged (day 7) worms. n = 46, 41, 43, 35, 45, 44, 42, 36; worms from 2 independent trials (left to right); mean ± SD; two-way ANOVA with Tukey’s multiple comparisons test. (G) TEM imaging of mitochondria in hypodermal cells of young (day 1) worms. Scale bar: 500 nm. (H) Quantification of mitochondrion size in young (day 1) worms. n = 23, 202, 49, 105 individual mitochondria from 1-2 images each from 2-4 worms (left to right); mean with 95% CI; one-way ANOVA with Sidak’s multiple comparisons test. (I) TEM imaging of mitochondria in hypodermal cells of aged (day 7) worms. Scale bar: 500 nm. (J) Quantification of mitochondrion size in aged (day 7) worms. n = 25, 109, 104, 136 individual mitochondria from 1-2 images each from 2-4 worms (left to right); mean with 95% CI; one-way ANOVA with Sidak’s multiple comparisons test. (G, I) Arrows indicate lipid droplets; arrowheads indicate mitochondria. ns p > 0.05; * p < 0.05; ** p < 0.01; *** p < 0.001; and **** p < 0.0001.

Overall we find that while the transcriptional and physical expansion of mitochondrial networks are tightly coupled in Complex I mutants, this coupling is lost when InsP3R function is impaired. Instead, InsP3R dysfunction exacerbates mitochondrial network expansion, while reducing biogenesis at the transcriptional level. The contrasting network expansion concomitant with downregulation of biogenesis transcripts may together suggest that the blunted transcriptional response arises through negative feedback in InsP3R/*itr-1*; Complex I/*nuo-6* mutants. This model provides a potential explanation for prior speculation from UPRmt screens that InsP3R signaling influences UPRmt outputs through indirect mechanisms^29,51–53^, but further studies are required. Here, the exacerbation of mitochondrial expansion despite reduced transcriptional biogenesis signatures suggests that post-transcriptional mechanisms, such as impaired turnover or structural remodeling machineries, are the primary drivers of both the mitochondrial and lifespan phenotypes downstream of InsP3R. Notably, the marked expansion of mitochondrial networks in Complex I/*nuo-6;* InsP3R/*itr-1* mutants occurs despite significantly reduced respiratory output (Fig. 2), consistent with a maladaptive response in which dysfunctional organelles accumulate in the absence of proper regulatory control.

### InsP3R control of mitochondrial damage-associated actin mediates lifespan extension in Complex I mutants

Based on the aberrant mitochondrial scaling and morphologies in InsP3R/*itr-1*; Complex I*/nuo-6* double-mutants (Figs. 3, 4), we next hypothesized that cytosolic calcium signaling might intersect with regulation of mitochondrial dynamics to play important roles in potentiating Complex I longevity. To begin dissecting potential roles for cytosolic mediators of calcium signaling, we first tested a role for calmodulin/*cmd-1*, which localizes to the cytosol and nucleus and directly binds Ca^2+^ released by the InsP3R to initiate diverse downstream signaling pathways. While calmodulin/*cmd-1* knockdown from hatch leads to developmental arrest, strikingly, we found that even post-developmental knockdown of calmodulin/*cmd-1* suppresses Complex I longevity (Fig. 5A), consistent with a role downstream of InsP3R/*itr-1*. This requirement for calmodulin coupled with the lack of an effect from *mcu-1* ablation strongly supports a model where cytosolic calcium signals potentiate mitochondrial-mediated longevity.

**Figure 5:**
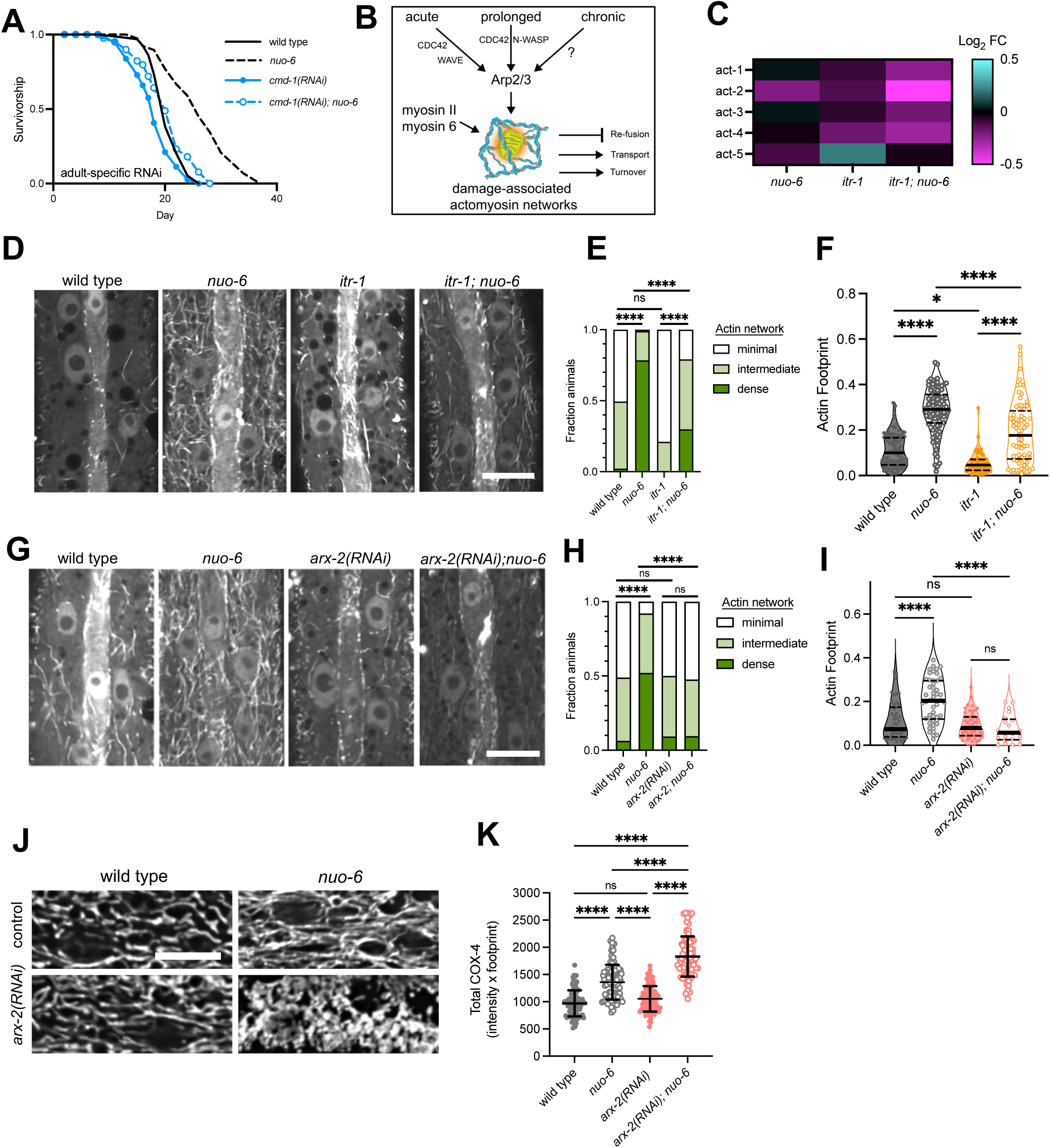
InsP3R signaling promotes actin remodeling in Complex I/*nuo-6* mutants to regulate mitochondrial dynamics. (A) Lifespan of Complex I/*nuo-6* mutants fed *cmd-1(RNAi)* from L4. n = 100 worms per condition. Statistical significance of lifespan curves was determined by a log-rank Mantel-Cox test. See Supplementary Table 1 for lifespan statistics. (B) Working model of mitochondrial stress-associated actin remodeling. (C) Heatmap comparing the expression of actin cytoskeletal mRNA transcripts in Complex I/*nuo-6*, InsP3R/*itr-1*, and InsP3R/*itr-1*; Complex I/*nuo-6* mutants. All genes normalized to WT. (D) Representative images of F-actin networks (LifeAct-GFP) in hypodermis of L4 worms. Scale bar: 10 µm. (E) Categorization of F-actin network density. n = 89, 102, 52, 77 worms from 4 independent trials (left to right); Kruskal-Wallis test with Dunn’s multiple comparisons test. (F) Quantification of actin footprint. n = 90, 98, 48, 76 worms from 4 independent trials (left to right); median with quartiles; one-way ANOVA with Tukey’s multiple comparisons test. (G) Representative images of F-actin networks (LifeAct-GFP) in hypodermis of L4 worms fed *arx-2(RNAi)*. Scale bar: 10 µm. (H) Categorization of F-actin network density in worms fed *arx-2(RNAi).* n = 47, 50, 54, 21 worms from 3 independent trials (left to right); Kruskal-Wallis test with Dunn’s multiple comparisons test. (I) Quantification of actin footprint in worms fed *arx-2(RNAi)*. n = 47, 50, 54, 21 worms from 3 independent trials (left to right); median with quartiles; one-way ANOVA with Tukey’s multiple comparisons test. (J) Representative images of mitochondrial networks (COX-4::eFGP) in hypodermis of young (day 1) worms fed *arx-2(RNAi).* Scale bar: 10 µm. (K) Quantification of total COX-4 (mean COX-4::eGFP intensity x mitochondrial footprint). n = 98, 87, 96, 84 worms from 4 independent trials (left to right); mean ± SD; one-way ANOVA with Sidak’s multiple comparisons test. ns p > 0.05; * p < 0.05; and **** p < 0.0001.

Calmodulin interacts with many proteins, so we next sought to test potential downstream targets of calmodulin more directly linked to mitochondrial dynamics. We focused first on calcium/calmodulin-dependent protein kinase II (CaMKII)/*unc-43* and calcineurin/*tax-6*, which are both linked to lifespan as well as regulation of mitochondrial fission through dynamin-related protein (DRP)-1^52^. While calcineurin/*tax-6* knockdown alone was sufficient to extend lifespan as previously shown^54^, Complex I/*nuo-6* dependent lifespan extension was independent of both *unc-43* and *tax-6* (Extended Data Fig. 4A,B). Though we observed no clear role for these potential upstream regulators of fission, we also attempted to determine more directly whether DRP-1 fission complexes are altered in InsP3R and Complex I mutants. We employed a GFP::DRP-1 line to visualize the formation of DRP-1 oligomers *in vivo*, but discovered fluorescence labeling of DRP-1 significantly impacts its function (Extended Data Fig. 4C), consistent with prior *in vitro* findings^54^. To mitigate this issue, we maintained *gfp::drp-1* as a heterozygote alongside a wild type *drp-1* copy across all backgrounds and used *nuo-6(RNAi)* in place of *nuo-6(qm200)* mutants to avoid synthetic phenotypes (Methods). With these caveats, we found that both the number and mean intensity of GFP::DRP-1 complexes were actually increased in the Complex I/*nuo-6(RNAi)* background, and to an even greater extent in InsP3R/itr-1; Complex I/*nuo-6(RNAi)* animals (Extended Data Fig. 4D-I). This inverse correlation between fused mitochondrial morphologies and the number of fission complexes across genotypes may suggest that fission complexes exist in a poised, but inactive or impaired state in the InsP3R/*itr-1*; Complex I/*nuo-6(RNAi)* condition. Notably, given that *nuo-6(RNAi)*-mediated longevity was independent of InsP3R/*itr-1,* we found that the mitochondrial hyperexpansion observed in InsP3R/*itr-1*; *nuo-6(qm200)* animals did not occur in InsP3R/*itr-1; nuo-6(RNAi)* conditions (Extended Data Fig. 4J,K). While technical challenges in our system limit our ability to explore how fission complexes are differentially regulated under these conditions, future studies with improved tools or alternative models will be useful in dissecting the molecular regulation of DRP-1 in relevant contexts.

An alternative pathway for regulating mitochondrial dynamics has emerged from recent *in vitro* studies, collectively revealing that Ca^2+^ signals play a role in triggering actin remodeling around dysfunctional mitochondria. These actin ‘cages’ can prevent fusion of damaged mitochondria within the network, enhance metabolic rewiring, and promote dispersal and trafficking of damaged mitochondria for turnover^31–34^ (Fig. 5B). Intriguingly, if these damage-induced actin networks are impaired during prolonged mitochondrial dysfunction, mitochondrial networks expand and individual mitochondria become enlarged or swollen^32^, similar to what we observe in InsP3R; Complex I mutants (Fig. 4). Consistent with a link to actin regulation, we noticed a trend of downregulation (14-43%) of 4 out of 5 actin genes arising in the double-mutants in our transcriptomic dataset (Fig. 5C), further suggesting the potential for altered actin regulation in our model. While roles for this type of actin remodeling are so far unexplored *in vivo* or in aging contexts, we reasoned that impaired InsP3R signaling during Complex I dysfunction could result in defective actin remodeling, thus explaining the larger and more hyperfused network we observe in double-mutants. To observe actin remodeling more directly, we employed a LifeAct::GFP reporter for F-actin and tested whether mitochondrial dysfunction causes similar actin remodeling in *C. elegans*. Indeed, we observed a roughly three-fold increase in F-actin networks in Complex I/*nuo-6* mutants (Fig. 5D-F), demonstrating pronounced actin remodeling in response to mitochondrial dysfunction in *C. elegans*. Furthermore, the increase in F-actin formation was impaired in InsP3R/*itr-1* mutants, consistent with its role in the early responses to mitochondrial depolarization *in vitro*^55^ (Fig. 5D-F). By contrast, we found that *nuo-6(RNAi)* does not induce significant actin network formation (Extended Data Fig. 4L), further supporting that this InsP3R–actin remodeling axis distinguishes the mutant from knockdown-based ETC perturbation. Together, these results reveal that mitochondrial damage-associated actin remodeling is conserved in *C. elegans* and promoted by InsP3R signaling during chronic mitochondrial dysfunction.

Recent work from multiple groups has delineated several factors that mediate damage-associated actin remodeling. The remodeling occurs in partially distinct acute versus prolonged phases of mitochondrial stress, and each phase may involve molecular mechanisms that are partly overlapping and partly unique^34,55^. Chronic mitochondrial dysfunction, for example in models of Leigh syndrome, also induces actin remodeling, though the regulation and consequences in these chronic contexts are not as well understood^31,32,55^. Importantly, the factor shown to be important across all phases of mitochondrial damage-associated actin remodeling is the nucleating and branching Arp2/3 complex^31,55^. We therefore tested whether Arp2/3 is also required for damage-associated F-actin polymerization in *C. elegans* during Complex I/*nuo-6* impairment. When we targeted Arp2/3 function via *arx-2(RNAi)*, we found that *arx-2* had little effect on basal F-actin networks, but fully suppressed the actin networks induced by Complex I/*nuo-6* dysfunction, confirming a key, conserved role for Arp2/3 in forming and/or stabilizing these stress-dependent actin networks (Fig. 5G-I). We also tested whether Arp2/3 plays a role in regulating mitochondrial networks in Complex I/*nuo-6* mutants by imaging mitochondrial COX-4::GFP markers. Again, Arp2/3/*arx-2* knockdown phenocopied InsP3R mutants by demonstrating an enhanced expansion of mitochondrial networks upon *arx-2* inhibition (Fig. 5J,K). Collectively these results support a model where InsP3R signaling and Arp2/3-dependent actin remodeling regulate stress-dependent mitochondrial dynamics in *C. elegans* as well as in mammalian models.

Next, if InsP3R/*itr-1* effects on lifespan were mediated via actin remodeling, we predicted that impairing damage-associated actin remodeling downstream of InsP3R signaling would phenocopy the suppression of Complex I/*nuo-6* associated longevity. We fed animals *arx-2* dsRNA, and while Arp2/3 impairment reduced lifespan compared to controls, this treatment completely suppressed Complex I-dependent lifespan extension (Fig. 6A). We also tested an alternative Arp2/3 subunit, *arx-4*, which yielded near-identical results (Fig. 6B). Overall, these findings demonstrate a novel role for the Arp2/3 complex and damage-associated actin remodeling more broadly in promoting longevity during mitochondrial stress contexts. While Arp2/3 activity is centrally important to damage-associated actin responses, we hypothesized that additional factors likely contribute to adaptive responses during chronic mitochondrial stress. Based on current models of acute damage-associated actin (ADA) and prolonged damage-associated actin (PDA) formation, we next focused on potential roles for additional actin nucleation pathways and actomyosin regulators known to influence mitochondrial dynamics during stress. In addition to Arp2/3 branched actin networks, some forms of damage-associated actin also involve formin-mediated linear filament assembly^55^. Indeed, we found that RNAi targeting of the formin FHOD-1 completely suppressed Complex I/*nuo-6* mutant lifespan extension (Fig. 6C), suggesting an important role for linear actin nucleation in our mitochondrial longevity model, potentially by promoting stable actin structures required for prolonged mitochondrial segregation or trafficking. Previously, Cdc42 and Wiskott-Aldrich syndrome protein (WASP)-family proteins were found to activate the nucleation by Arp2/3 and formins in mitochondrial damage paradigms^33,55^, so we tested roles for these genes in mediating mitochondrial longevity. We found that knockdown of both *cdc-42* and *wsp-1* largely suppressed lifespan extension of Complex I mutants from 45% to 37.5% and 45% to 20% (Fig. 6D,E). Finally, myosins cooperate with actin networks to perform diverse roles in mitochondrial dynamics, including actin-based scaffolding and constriction of mitochondria for fission, stabilization of damage-associated actin networks, and disassembly of ‘aggregated’ mitochondria for delivery to autophagosomes following damage^56^. Notably, some unconventional myosins associated with damage-associated actin subpopulations also use calmodulin/CMD-1 itself as a regulatory light chain^57^. While genetic redundancy among non-muscle myosins makes genetic dissection of specific myosin genes challenging in this system, these functions are often regulated by upstream phosphorylation of myosin regulatory light chains via myosin light chain kinase (MLCK), a well-established target of calmodulin. In contrast to CaMKII and calcineurin, we found that RNAi against MLCK-1 substantially suppressed Complex I/*nuo-6* lifespan extension (Fig. 6F), highlighting a role for calmodulin-mediated actomyosin signaling in mitochondrial stress adaptation. Further, MLCK-1 appears to play no role in lifespan extension downstream of insulin signaling pathways (Extended Data Fig. 5), indicating specificity to mitochondrial longevity paradigms. Overall, our findings integrate emerging *in vitro* models with *in vivo* genetic evidence to support a mechanism in which InsP3R-derived Ca²⁺ signals act through calmodulin and MLCK to coordinate actomyosin-based scaffolding and remodeling of mitochondrial networks, thereby facilitating mitochondrial quality control and enabling lifespan extension during chronic mitochondrial stress.

**Figure 6:**
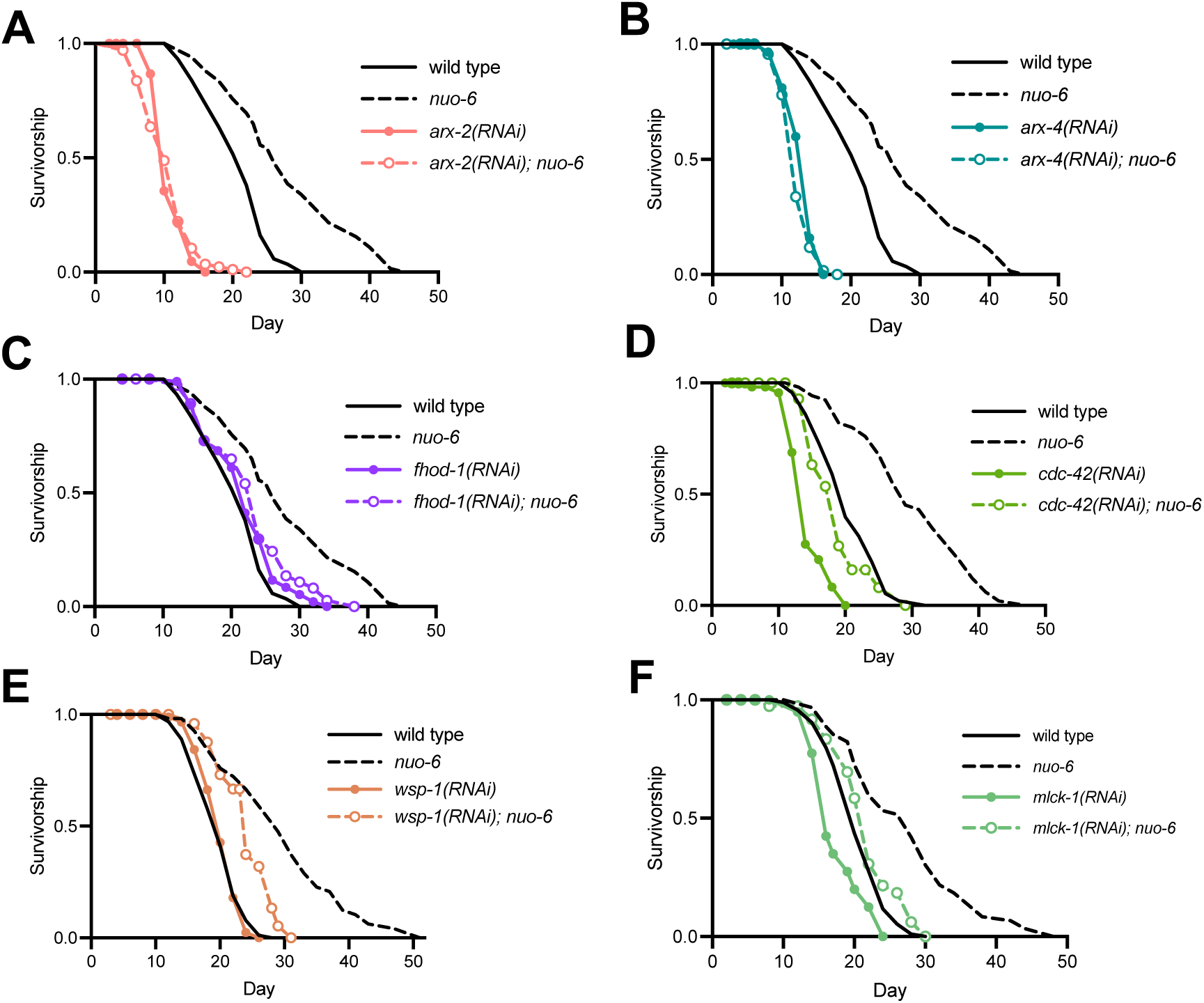
An actomyosin regulatory network is required for lifespan extension during Complex I impairment. (A) Lifespan of Complex I/*nuo-6* mutants fed *arx-2(RNAi).* n = 97-100 worms per condition. (B) Lifespan of Complex I/*nuo-6* mutants fed *arx-4(RNAi).* n = 97-100 worms per condition. (C) Lifespan of Complex I/*nuo-6* mutants fed *fhod-1(RNAi).* n = 97-100 worms per condition. (D) Lifespan of Complex I/*nuo-6* mutants fed *cdc-42(RNAi).* n = 67-100 worms per condition. (E) Lifespan of Complex I/*nuo-6* mutants fed *wsp-1(RNAi).* n = 80-105 worms per condition. (F) Lifespan of Complex I/*nuo-6* mutants fed *mlck-1(RNAi).* n = 100 worms per condition. Statistical significance of lifespan curves was determined by a log-rank Mantel-Cox test. See Supplementary Table 1 for lifespan statistics.

Finally, because mitochondrial damage-induced actomyosin remodeling collectively shifts the network balance towards fragmentation by promoting DRP-1 fission^57^ and by inhibiting re-fusion^32^, we hypothesized that restoring the ability of mitochondrial networks to fragment would rescue lifespan extension in InsP3R/*itr-1* mutants. To test this we inhibited the outer mitochondrial membrane fusion factor Mfn/*fzo-1*, revealing robust mitochondrial fragmentation across all genotypes (Fig. 7A,B). Furthermore, Mfn/*fzo-1* impairment uniquely reverses the mitochondrial hyperexpansion and elevated COX-4 accumulation in double-mutants, restoring both footprint and COX-4 levels to those observed in Complex I/*nuo-6* single mutants (Fig. 7C & Extended Data Fig. 6A,B). Strikingly, Mfn/*fzo-1(RNAi)* also restored lifespan extension in Complex I/*nuo-6*; InsP3R/*itr-1* double mutants (Fig. 7D,E), indicating that reversing outer membrane fusion is sufficient to bypass defects in stress adaptation in InsP3R mutants. Neither inhibition of mitochondrial fission (*drp-1(RNAi)*) nor disruption of inner membrane fusion (Opa1/*eat-3(RNAi)*) produces this rescue effect (Extended Data Fig. 6C,D).

**Figure 7:**
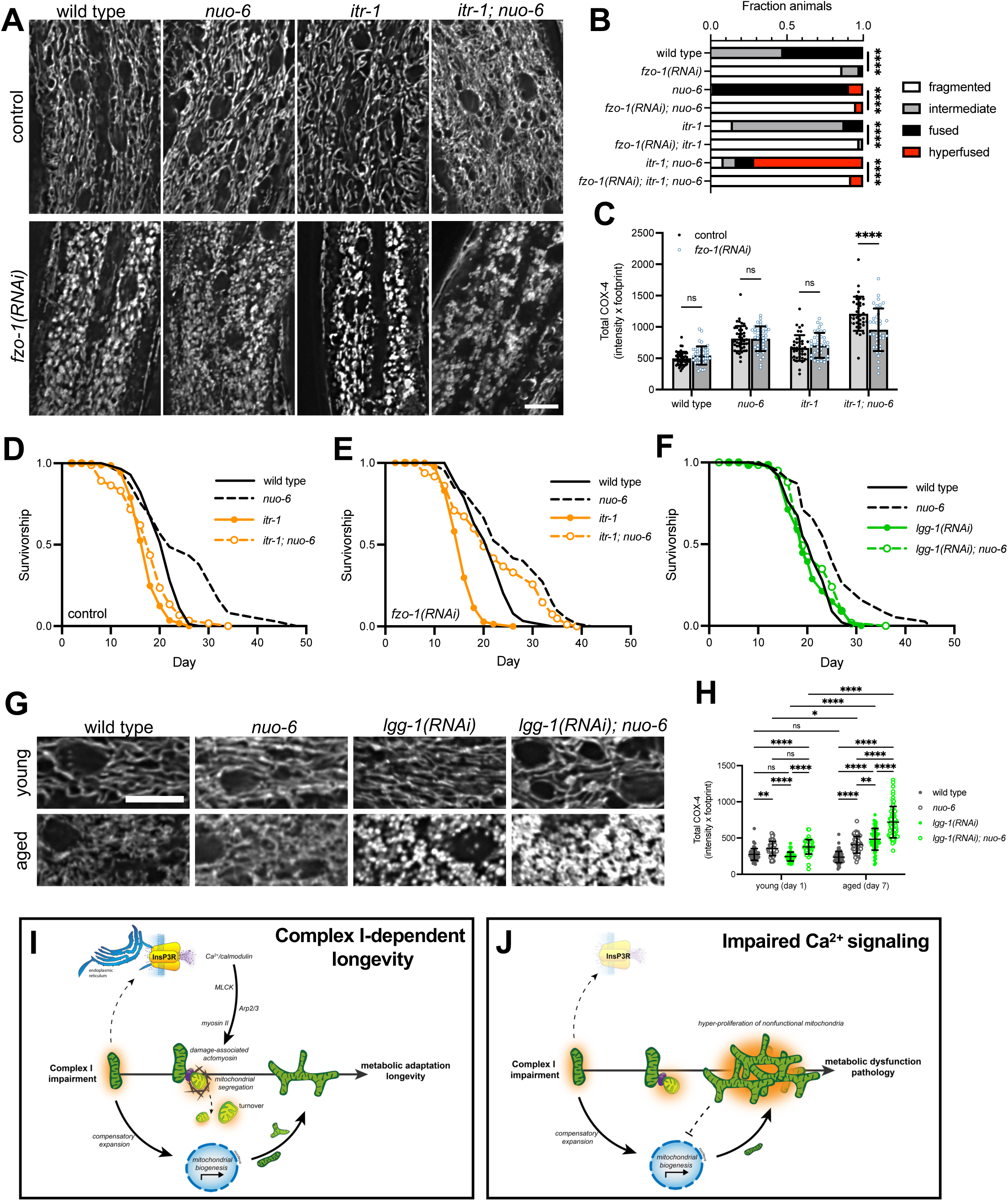
Driving mitochondrial fission rescues longevity in InsP3R mutants and autophagy impairment mimics InsP3R/*itr-1* loss. (A) Representative images of mitochondrial networks (COX-4::eGFP) in hypodermis of young (day 1) worms fed control or *fzo-1(RNAi).* Scale bar: 10 µm. (B) Categorization of mitochondrial morphologies in worms fed control or *fzo-1(RNAi).* n = 44, 44, 43, 44, 41, 40, 35, 38 worms from 2 independent trials (top to bottom); Kruskal-Wallis test with Dunn’s multiple comparisons test. (C) Quantification of total COX-4 (mean COX-4::eGFP intensity x mitochondrial footprint). n = 43, 44, 42, 43, 41, 40, 35, 38 worms from 2 independent trials (left to right); mean ± SD; two-way ANOVA with Sidak’s multiple comparisons test. (D, E) Lifespan of InsP3R/*itr-1* and Complex I/*nuo-6* double mutants fed control (D) and *fzo-1(RNAi)* (E). n = 100 worms per condition. (F) Lifespan analysis of Complex I*/nuo-6* mutants fed *lgg-1(RNAi).* n = 100 worms per condition. (G) Representative images of mitochondrial networks (COX-4::eGFP) in hypodermis of young (day 1) and aged (day 7) worms fed control or *lgg-1(RNAi).* Scale bar: 10 µm. (H) Quantification of total COX-4 (mean COX-4::eGFP intensity x mitochondrial footprint). n = 63, 50, 54, 53, 74, 66, 62, 61 worms from 4 independent trials (left to right); mean ± SD; two-way ANOVA with Tukey’s multiple comparisons test. (I, J) Model for how the InsP3R regulates mitochondrial dynamics during Complex I impairment (I) and the effect of impaired Ca^2+^ signaling during Complex I impairment (J). Statistical significance of lifespan curves was determined by a log-rank Mantel-Cox test. See Supplementary Table 1 for lifespan statistics. ns p > 0.05; * p < 0.05; ** p < 0.01; and **** p < 0.0001.

Given the diverse functional consequences of altered mitochondrial morphology, multiple mechanisms likely contribute to the lifespan rescue following Mfn/*fzo-1* impairment, including potential alterations in damage segregation, calcium and lipid handling, metabolic and redox signaling, and/or mitochondrial transport^28^. However, prior work has a key outcome of PDA-like actin remodeling is mitophagic turnover^58,59^, while elongated mitochondrial networks resist autophagic clearance^23,60^. These observations, combined with our findings that InsP3R signaling both promotes actin remodeling and constrains mitochondrial expansion, led us to hypothesize that InsP3R signaling prevents excessive accumulation of dysfunctional mitochondria by facilitating autophagic clearance. Supporting this hypothesis, we impaired autophagy via LC3/*lgg-1(RNAi)* and found that autophagy is indeed required for Complex I/*nuo-6*–mediated lifespan extension (Fig. 7F). Furthermore, impairing autophagy in Complex I/*nuo-6* mutants phenocopies the mitochondrial hyperexpansion (Fig. 7G,H) seen when both InsP3R/*itr-1* and Arp2/3 are impaired (Fig. 4A-F & Fig. 5J,K). Together these results indicate that the ability to limit the compensatory expansion of mitochondrial networks during chronic Complex I dysfunction is essential for longevity. By establishing that InsP3R signaling coordinates cytoskeletal remodeling with mitochondrial clearance, our findings define a conserved, calcium-dependent mechanism of mitochondrial quality control critical for organismal lifespan extension.

## Discussion

Here we uncover a critical new role for the InsP3R, and calcium signaling more broadly, in mediating lifespan extension during Complex I impairment. Specifically, our findings support a model in which cytosolic Ca^2+^ signals from the InsP3R coordinate mitochondrial dynamics through calmodulin-dependent actomyosin remodeling, thereby maintaining mitochondrial homeostasis and permissive conditions for longevity (Fig. 7I,J). Our study focuses on the consequences of impaired calcium signaling in contexts of life-extending ETC mutations, but cellular Ca^2+^ defects also commonly co-occur with mitochondrial dysfunction during advanced age and disease^25,40,61,62^. Thus, our findings suggest that age-associated disruptions in mitochondrial function and ER calcium dynamics can trigger a pathological vicious cycle, impairing quality control and driving greater mitochondrial dysfunction and disease. In this way, our work illuminates inter-organelle calcium dynamics and actin remodeling as key processes that differentiate beneficial mitochondrial stress from dysfunction, an area where the field has struggled to move beyond simple ROS- and UPRmt-based models. Along these lines, our accompanying transcriptomic resource provides a foundation for exploring additional roles for calcium signals in shaping cellular and organismal responses to mitochondrial dysfunction and aging.

One of the most striking advances of this study is the demonstration that mitochondrial stress-dependent actomyosin remodeling, which had previously been defined *in vitro*, plays an essential role in promoting mitochondrial homeostasis and longevity in an animal model of chronic ETC impairment. Specifically, we show that Arp2/3, formin FHOD-1, and upstream regulators CDC-42 and WSP-1, are required for Complex I-induced longevity, placing InsP3R-calmodulin signaling upstream of actomyosin dynamics in this pathway. These findings reveal a conserved calcium-cytoskeleton-mitochondria axis with key roles in aging and lifespan. Identifying specific myosin mediators and understanding how these actin subpopulations intersect with mitochondrial fission/fusion, mitophagy and transport machineries in these chronic contexts remain important questions for future studies. Finally, our findings establish the regulation of mitochondrial expansion as a critical factor distinguishing adaptive stress responses from maladaptive outcomes. An important next step will be to determine how unchecked mitochondrial expansion promotes cellular dysfunction. Potential explanations include enhanced propagation of ROS, disruption of inter-organelle interactions and communication, or simply draining metabolic resources towards futile maintenance of irreparable organelles.

Our finding that the canonical InsP3R-MCU pathway is dispensable for Complex I-mediated lifespan extension reinforces the importance of extra-matrix signaling pathways in mitochondrial homeostasis, complementing prevailing models that emphasize ER-to-matrix Ca^2+^ flux^63,64^. Our data from *mcu-1* mutant worms are consistent with the relatively minor phenotypes associated with MCU knockout mice^41,55,65–68^, and support a model where extra-matrix Ca^2+^ signaling pathways and redox shuttles may play larger roles in mitochondrial metabolism and homeostasis than previously anticipated^41,55,65–68^. In our model, reduced membrane potential in Complex I mutants likely limits matrix Ca^2+^ uptake. This impaired Ca^2+^ uptake during Complex I dysfunction provides a potentially simple mechanism by which InsP3R activity may respond to mitochondrial stress, as previous work showed that mitochondrial Ca^2+^ uptake suppresses InsP3R activation by removing positive feedback by cytosolic Ca^2+^ ions^2,3,42^. In the future, it will also be useful to explore whether Ca^2+^-regulated redox pathways, such as the glycerol phosphate or malate-aspartate shuttles, are important for InsP3R-dependent control of bioenergetics and stress adaptation in long-lived mitochondrial mutants.

While early models suggested a unified mechanism promoting longevity during mitochondrial stress^69^, mounting evidence points to mechanistic heterogeneity across different ETC perturbations. For example, a recent suppressor screen showed that virtually all compensatory responses to a mutation in Complex I acted locally within the affected complex itself^69^, underscoring the specificity of these adaptations. Likewise, distinct modes of perturbation can engage parallel pathways: interventions that extend lifespan through stable ETC mutations are additive when combined with RNAi-induced ETC disruption^42^. Consistent with this, we show that InsP3R function is required for longevity in *nuo-6* mutants but dispensable for *nuo-6* knockdown, revealing a striking example of genetic divergence based on how Complex I is perturbed. Moreover, while InsP3R/*itr-1* mutants tolerate Complex I knockdown, they exhibit profound developmental arrest in response to ETC inhibition at Complexes III, IV, or V, indicating differential requirements for InsP3R function when distinct Complexes are perturbed. This difference may reflect distinct physiological demands. Specifically, Complex I mutation allows alternative electron entry points into the ETC, limiting the need for redox compensation, while impairments in Complexes III, IV and V present more severe energetic bottlenecks that increase reliance on potential InsP3R roles in regulating redox balance. Collectively, these findings reveal InsP3R signaling plays important, context-specific roles in adapting to ETC dysfunction. More broadly, these results suggest differences in how ETC perturbations impact calcium signaling and cytoskeletal remodeling could underlie the mechanistic diversity in mitochondrial longevity paradigms.

Overall, our work provides a framework for understanding how organelle-scale calcium dynamics intersect with cytoskeletal remodeling to mediate stress adaptation and lifespan regulation. Our findings reveal previously unknown roles for InsP3R-dependent calcium signaling pathways in promoting lifespan extension under chronic mitochondrial stress. These results broaden the scope by which ER calcium signaling is promoting mitochondrial homeostasis, expanding from the canonical ER-to-matrix Ca^2+^ flux model and highlighting critical roles for cytosolic signaling, cytoskeletal remodeling and autophagy-mediated turnover. Given the widespread association between impaired ER-mitochondrial communication, cytoskeletal dysfunction, and aging-related diseases, the pathways defined here offer promising therapeutic targets for promoting mitochondrial health and longevity.

## Methods

### *C. elegans* strains and husbandry

See Supplementary Table 3 for complete list of strains used in this study. Worms were grown and maintained at 20 °C on standard nematode growth media (NGM) seeded with *E. coli* (OP50-1). *E. coli* cultures were grown from single colonies overnight in LB at 37 °C. 100 µL of liquid culture was then seeded onto NGM plates and bacterial lawns were allowed to grow for 2 days at room temperature before use.

### RNAi experiments

RNAi experiments were conducted by feeding worms sequence confirmed RNAi clones (*E. coli* - HT115) from the Ahringer RNAi library (Source Bioscience)—see Supplementary Table 4 for RNAi sequences. Experiments utilizing RNAi were performed using LB cultures and NGM plates supplemented with 100 µg/mL carbenicillin. RNAi was induced using 100 µL 0.1 M IPTG (supplemented with 100 µg/mL carbenicillin and 12.5 µg/mL tetracycline) at least 1 hour before worms were added to plates. *atp-3(RNAi)* was diluted 1 in 10 (10%) to measure development of worms at the maximal lifespan extension of complex V as previously noted^70^. *cmd-1(RNAi)* treatment was activated starting at L4 to avoid developmental arrest. All other RNAi experiments were treated from hatching.

### Developmental Assay

Worms were synchronized by timed egg lay at 20 °C on NGM plates supplemented with carbenicillin and then ∼50 eggs per condition were transferred to activated RNAi plates (time = 0). Worms were inspected and scored for 10 days following egg-lay, and vulval development was used to distinguish between worms arrested at late larval stages and sterile adults. Unhatched eggs and desiccated worms were censored. Four independent replicates were performed.

### InsP3 Sponge Construction

To generate an InsP3 sponge system, we adapted the design of Walker *et al*.^37^ with minor modifications. To create plasmid pGF1 (*eft-3p::supersponge::AID::mKate2::unc54 3’UTR*) we digested pDB6 using XmaI/NheI to create a backbone with 5’ *eft-3p* and 3’ *mKate2* sequences. The InsP3 sponge was generated by cloning the sequence representing amino acids M254 to K670 from *itr-1d* cDNA using forward primer: GACCCGGGATGTTTTTGCTTTTTGATG and reverse primer: CTGGATCTTTAGGCTTTGGATTATTGTGC. cDNA was derived from *itr-1(sy290)* allele, which contains the mutated InsP3 binding site that doubles affinity for InsP3^37^. Auxin-inducible degron (AID) sequence was amplified from pLZ29 (*eft-3p::AID::EmGFP*) using forward primer: ACAATAATCCAAAGCCTAAAGATCCAGCC and reverse primer: TTAGCTAGCACTCTTCACGAACGCC. The InsP3 binding domain was fused with AID via PCR using the IP3 binding domain forward primer (containing XmaI site) and AID reverse primer (containing NheI site). The resultant InsP3 supersponge::AID sequence was then ligated into the backbone. This plasmid was microinjected into N2 animals to produce *bugEx7[eft-3p::IP3::supersponge::AID::mKate2]* and subsequently gamma irradiated and outcrossed 3x to produce integrated strain BUZ117.

### Confocal Fluorescence Microscopy

Synchronized worms at indicated ages were picked onto a 10% agarose pad with ∼3 µL 0.1 µm Polybead microsphere suspension (Polyscience) for immobilization and then a cover slip (VWR) was added on top to mount worms^72^. Confocal microscopy was performed on a Nikon Eclipse Ti2 with CSU-W1 spinning disk and Plan-Apochromat 100x/1.49 objective. Mitochondrial targeted GCaMP7b, COX-4::eGFP, LifeAct-GFP, and GFP::DRP-1 were imaged by 488 nm laser excitation and ET525/36m emission filter. Mitochondrial targeted mKate2 and TMRE were imaged with 561 nm laser excitation and ET605/52m emission filter. DIC images were also taken alongside the fluorescent images. Nikon NIS-Elements Advanced Research (NIS-AR) was used for image processing and analysis.

### Calcium imaging

Images of mitochondrial targeted GCaMP7b and mKate2 were taken in the anterior-most intestinal cells. Mitochondrial targeted GCaMP7b was imaged with 2X integration to increase signal strength. Mitochondrial networks were identified and masked via setting threshold of mKate2 signal after rolling ball background subtraction. Samples were blinded and thresholding parameters were manually adjusted for every image. Regions of interest (ROIs) were manually drawn to encompass a single anterior intestine cell. Subsequently, the mean intensity of GCaMP7b and mKate2 were measured in the entire mitochondrial network of single intestinal cells, and then the ratio of GCaMP7b:mKate2 signal was calculated.

### Mitochondrial imaging

Endogenously tagged COX-4::eGFP was imaged in the anterior intestine and hypodermis of day 1 and day 7 worms. Deconvolution and denoising were applied to get sharper images. ROIs were manually drawn to include a single intestine cell in the anterior-most intestine or the hypodermis. In aged intestine images, bright spot detection (Nikon Elements) was used to exclude auto-fluorescent gut granules. Mitochondrial networks were identified and segmented by setting threshold via COX-4::eGFP signals. To objectively set threshold parameters across all groups, files were blinded and lower thresholding parameters were manually adjusted for every image. Within each ROI, the mean COX-4::eGFP intensity of segmented mitochondria, the area of segmented mitochondria, and ROI area were measured. Mitochondrial footprint (total segmented mitochondrial area divided by ROI area) and total COX-4 (mitochondrial footprint multiplied by mean COX-4::eGFP intensity) were calculated. To quantify mitochondrial networks based on their overall morphology, blinded image files were used to manually classify networks into one of four categories: fragmented, intermediate, fused, or hyperfused. Neurons, nuclei, and seam cells were omitted from ROIs.

### Actin network imaging

LifeAct-GFP was used to visualize filamentous actin in the hypodermis of mid-L4 worms. Worms were synchronized to mid-L4 by identifying the “christmas tree” stage of vulva development. All images were taken using 2X integration in the hypodermis just anterior to the vulva in a plane with visible nuclei. To enhance visualization of the actin networks, all images were denoised and sharpened (Nikon Elements). All files were blinded prior to analysis. ROIs were manually drawn to include the hypodermis while omitting seam cells and nuclei. Actin filament networks were segmented by manually setting threshold parameters and actin footprint (total segmented actin area divided by ROI area) was calculated. Blinded image files were also used to manually classify actin networks into one of three categories: dense, intermediate, or minimal.

### DRP-1 imaging, processing, and analysis

We encountered multiple technical challenges in establishing lines for the study of DRP-1 interactions with mitochondria. First, we discovered that animals homozygous for GFP-labeled DRP-1 exhibit grossly aberrant mitochondrial morphology, as suggested also in tissue culture studies^73^. However, animals heterozygous for DRP-1::GFP with one wild type copy exhibit mitochondrial morphology similar to controls (Extended Data Fig. 4C), leading us to adopt this heterozygous approach across all backgrounds. Secondly, we were unable to generate the 4-locus strain (*itr-1; nuo-6; GFP::drp-1;* mito-mKate2) ideally needed to image DRP-1-mitochondrial interactions. We therefore used RNAi to knockdown *nuo-6* in *itr-1; drp-1::GFP(+/-); mito-mKate2* animals. Egg lays were performed on activated RNAi plates (empty vector vs *nuo-6*). At L4 stage, worms were sorted under a fluorescence stereoscope to include only those heterozygous for the *GFP::drp-1* transgene. These worms were maintained on the corresponding RNAi lawns. At days 1 and 7, 20-30 worms for each condition were removed and mounted for imaging as described above. All images were taken around the middle portion of the anterior hypodermis. Because of fluorophore crosstalk, spectral unmixing was performed (Nikon Elements AR) with spectra for “eGFP” and “mTangerine” providing the cleanest separation of signal. Unmixed images were then subjected to automated 2-D deconvolution (green ex/em 488/535; red ex/em 561/620) and denoising (Denoise.ai module). Rolling-ball background removal was performed on each channel with identical settings for all images in the experimental replicate. ROIs were drawn around hypodermis cell hyp7, excluding seam cells, neurons, and unfocused regions. Mitochondria and DRP-1 puncta were segmented via thresholding and brightspot detection, respectively. Within each ROI, the total number of DRP-1 puncta was counted and normalized to the total area of segmented mitochondria. Mean DRP-1 puncta intensity was measured collectively across all puncta within the ROI.

### Respiration Analysis

OCR measurements were conducted on synchronized day 1 adults using Agilent Seahorse XFe96 flux analyzer and Wave desktop software according to the manufacturer’s instructions and adapted for *C. elegans*^45^. The heater of XFe96 flux analyzer was turned off before use and experiments were performed at 20-25 °C. The XFe96 sensor cartridge was hydrated with sterile water overnight at 37 °C without CO_2_ and then hydrated with seahorse XFe96 calibrant for a few hours before use. Worms were washed off plates and washed three times with M9 buffer. ∼20 worms were loaded to each well of cell culture microplates (Agilent Seahorse XF96) except the wells set for background correction. After calibration and equilibration, basal OCR was measured for 5 loops and each loop contains mixing for 2 minutes, waiting for 0.5 minutes and measuring for 2 minutes. 10 µM FCCP (final concentration) was injected to induce maximal oxygen consumption rate for 9 loops. 40 mM sodium azide (final concentration) was injected to inhibit mitochondrial respiration and non-mitochondrial OCR was measured for 4 loops. The precise worm numbers for each well were counted after experiments and raw OCR readings were divided by worm numbers to calculate the OCR per worm (pmol/minute per worm). The well was excluded if its raw OCR was over 600 due to high worm number. To diminish the impact of transient noise arising from worm movements and allow time for FCCP to fully equilibrate in the animals, basal OCR was calculated by averaging the OCR across measurement loops 2-5 and the maximal OCR across loops 11-14. Each condition had at least 6 wells for technical replicates in one experiment and at least three independent experiments were performed.

### Lifespan Analysis

All lifespan experiments were performed at 20 °C. Timed egg lays or hypochlorite bleaching were used to obtain synchronized populations. One day before adulthood (day 0), at least 100 mid-L4 worms were transferred to 5 or 10 fresh plates at a density of 10 or 20 worms per plate. To separate from progeny and avoid starvation, worms were transferred to fresh plates every day or every other day until the first deaths (∼10-12 days). Survival was scored every 1-2 days and worms were marked dead when they failed to respond to three taps on the head, tail, and middle body part. Worms were censored when they crawled off the plate and desiccated, lost vulval integrity during reproduction, or when embryos hatched internally. Log-rank Mantel-Cox tests were performed to determine statistical significance between two conditions.

### Lifespan analysis on solid dietary restriction (DR) plates

Solid dietary restriction assays were performed as previously described^74^ with the following modifications. Plates were prepared with a bacterial concentration of 10^11^ cfu/mL for *ad libitum* (AL) and 10^8^ cfu/mL for DR. After diluting bacteria to the correct concentrations in LB, the diluted bacterial stocks were put in a shaking incubator at 37 °C for 1 hour. After shaking for 1 hour, 50 mg/mL kanamycin and 100 mg/mL carbenicillin were diluted and mixed into the diluted bacterial stocks at 1:11 ratio prior to seeding plates. All plates were prepared in advance and stored at 4 °C before use. Worms were synchronized with timed egg lays on standard NGM plates with OP50-1 grown using standard techniques. For each condition, 150 day 1 adults were transferred to 15 AL or DR plates at 10 worms per plate.

### RNA-Seq Analysis

Experiments were performed with three biological replicates. About 1000 worms were synchronized by hypochlorite bleaching and grown to L4 on NGM plates seeded with HT115 *E. coli*. Worms were collected and washed three times with M9 buffer to remove bacteria. After the final wash, pelleted worms were resuspended in Qiazol, snap frozen in liquid nitrogen, and stored at –80 °C until RNA extraction. For RNA extraction, worm samples were disrupted in three freeze thaw cycles with Qiazol between 37 °C water bath and snap freezing in liquid nitrogen. Then samples were incubated and vortexed with chloroform at room temperature to extract RNA. RNA was purified through Qiagen RNeasy mini kit per manufacturer instructions and treated with Qiagen RNase-Free DNase on the column to get rid of DNA contamination.

RNA quality analysis, cDNA library, and Next-Generation sequencing were performed by VUMC Vanderbilt Technologies for Advanced Genomics (VANTAGE). RNA quality was analyzed (BioAnalyzer) with all samples having an RNA integrity number of 10. cDNA library was constructed with stranded mRNA (polyA-selected) library preparation kit (NEB). Paired-end 150 bp sequencing was performed on the Illumina NovaSeq6000 targeting 50M reads per sample. Analysis was performed through the Illumina Dragen pipeline. Quality control analysis via MultiQC indicated high quality reads with mapping rates >96% over >75 million total reads per replicate. Alignment to the genome (UCSC ce10) was performed via STAR^73^; gene counts and normalization were generated through Salmon (https://combine-lab.github.io/salmon/); and the DESeq2 algorithm (https://bioconductor.org/packages/release/bioc/html/DESeq2.html). The list of DE genes from each genotype relative to wild type were then trimmed to remove genes with 0 reads or status = LOW (< 10 transcripts per kilobase million).

To process for enrichment analysis, cutoffs for DE genes were set as >1.5-fold change and P_adj_ < 0.01, and these subsets of genes were contrasted across genotypes. WormCat 2.0^45^ (http://www.wormcat.com/) was used to analyze gene set enrichment using annotations from the whole genome (v2). Terms from WormCat annotation “Category 2” were used to identify enriched cellular processes.

### qRT-PCR

Synchronized worm populations were grown on RNAi from hatching and collected at L2. For RNA extraction, worms were resuspended in Qiazol then treated to three freeze-thaw cycles alternating between 37 °C and liquid nitrogen. Samples were then treated with chloroform and the RNA containing phase was collected and purified using a Qiagen RNeasy mini kit. cDNA was generated using the SuperScript VILO cDNA synthesis kit per manufacturer’s instructions. TaqMan qRT-PCR was run on a Bio-Rad CFX96 Thermal Cycler. Normalized fold change was calculated following the 2^-ΔΔCT method. The following TaqMan gene expression assays were used: *iscu-1* (Ce02467253_g1) as the internal control, *cco-1/cox-5b* (Ce02412619_g1), and *atp-3* (Ce02417729_g1).

### TMRE staining and imaging

500 uL of a 0.1 mM TMRE solution (0.1 M TMRE stock in DMSO diluted 1:1000 in M9) was added to an NGM plate pre-seeded with OP50-1. Plates were allowed to dry for ∼8 hrs, then L4 worms were picked onto TMRE plates and fed overnight. After 16 hours, worms were picked to standard NGM plates with OP50-1 for 1 hour to remove excessive dye before imaging. TMRE solution, plates, and treated worms were protected from light before imaging. TMRE images were taken in the hypodermis of day 1 adults. Deconvolution and rolling ball background subtraction were applied to get sharper images; brightspot detection was used to exclude TMRE aggregates. Prior to analysis all image files were blinded. Mitochondria were segmented by manually setting the threshold. ROIs were manually drawn to include the hypodermis and within the ROI the mean intensity of the segmented mitochondria was measured.

### Western Blotting

Synchronized L4/young adults were collected, washed three times with M9 buffer, and snap frozen in liquid nitrogen. Three independent biological replicates of worm samples were collected. Identical volumes of RIPA buffer (50 mM Tris, 150 mM NaCl, 1 mM EDTA, 1% Triton X-100, 0.1% sodium deoxycholate, 0.1% SDS, pH 7.5) supplemented with protease inhibitors and phosphatase inhibitors were added to worm samples and then samples were lysed by sonication. Lysates were centrifuged at 16,000 g for 15 minutes at 4 °C and supernatant protein concentration was measured using a Pierce BCA protein assay kit. An equal amount of protein was heated at 95 °C for 15 minutes with 4x Laemmli Sample Buffer added. Samples containing 40-60 µg protein were loaded to a gel and once separated, proteins were wetly transferred to a PVDF transfer membrane. The PVDF membrane was blocked with 5% milk in TBST or 5% BSA-TBS. Following incubation with the primary antibody (For p-PDH: anti-p-PDH (1:1000), anti-beta-actin (1:2000); For total PDH: anti-PDH (1:1000), anti-tubulin (1:5000)) the PVDF membrane was washed three times for 10 minutes each with TBST and then incubated with the secondary antibody (For p-PDH: HRP-conjugated anti-rabbit (1:2000), HRP-conjugated anti-mouse (1:10,000); For total PDH: AF488 anti-rabbit (1:10,000), IRDye 680 anti-mouse (1:10,000)). To detect HRP-conjugated antibodies, signals were developed using Pierce ECL Western Blotting Substrate, imaged with an Amersham Imager 600, and bands were quantified using ImageJ. Fluorescently labeled antibodies were imaged with an Odyssey Imaging System and bands were quantified using Image Studio.

### Transmission Electron Microscopy (TEM)

Worms were transferred to a conical with 5 mL of M9 buffer and spun down at 2000 rpm three times to wash bacteria off. Excess supernatant was removed. A pellet of live *C. elegans* was resuspended in 0.15 M sucrose prepared in M9 buffer and loaded into 200 µm deep well of a A-type carrier (Leica). The assembly was covered with flat side of B-type carrier (Leica) and vitrified using high-pressure freezing machine (Leica EM ICE). The frozen specimens were stored in liquid nitrogen until further processing. The freeze-substitution (FS) process was adapted from Bélanger *et al.*^74^. FS was performed using a cocktail of 0.5% glutaraldehyde and 0.1% tannic acid in acetone. Samples were transferred into automatic freeze-substitution machine (AFS2, Leica) pre-cooled to -140 °C and warmed to -90 °C over a period of 3 hours where they were then kept for 114 hours. At the end of a hold period, samples were washed 4 times for 30 minutes each with acetone. The final acetone wash was replaced with 2% osmium tetroxide in acetone. Following this, samples were warmed to -20 °C over a period of 12 hours after which temperature was maintained at -20 °C for an additional 10 hours before further warming to 0 °C over a period of 4 hours. Samples were then transferred into an ice filled bath and washed 4 times for 30 minutes each with acetone and infiltrated with Spurr’s resin (Electron Microscopy Sciences). During final steps of resin infiltration, individual worms were transferred from carriers into resin molds with a fine needle and samples were cured in an oven at 60 °C for 72 hours. Thin sections (70 nm) were cut using a Leica UC7 and a Diatome diamond knife. Thin sections were positioned on grids and stained with uranyl acetate and Reynold’s lead citrate. These sections were then imaged on a Tecani T12 TEM operating at 100 keV using an AMT CMOS camera. Mitochondrial area and footprint were quantified using Amira software. A mask of the mitochondria and hypodermis were manually drawn using the freehand masking tool in Amira, then area was measured.

### Quantification and Statistical Analysis

GraphPad Prism (versions 9 & 10) was used for statistical analysis and plotting graphs. Comparisons between two groups were determined by two-tailed unpaired student’s t test. For multiple comparisons, one- or two-way ANOVAs were performed with post-hoc tests. Two-way ANOVAs were used to compare multiple genotypes between young and old ages. Data for mitochondrion size was distributed on a log normal scale. Confocal and TEM image analyses were performed on blinded files. Statistical tests, p-values, sample size, and error bars are indicated in figure legends.

## Data Availability

All source data are provided with this paper. RNA-seq data are available through GEO: GSE297429. Worm strains generated in this study will be available on request and/or deposited to the Caenorhabditis Genetics Center (CGC) repository as appropriate. This paper does not report original code. Any additional information required to reanalyze the data reported in this paper is available upon request. Further information and requests for data, resources, and reagents should be directed to and will be fulfilled by the lead contact, Kristopher Burkewitz.

## Acknowledgements

We thank Vivian Gama, Maulik Patel and all members of the Burkewitz lab for discussion and feedback related to this manuscript. We would also like to thank the Patel lab for generously providing us with reagents and Suhong Xu for worm strain SHX377. We would like to acknowledge the Caenorhabditis Genetics Center (NIH Office of Research Infrastructure Programs P40 OD010440) for providing some worm strains used in this work. We thank the Washington University Center for Cellular Imaging (WUCCI) at the Washington University School of Medicine for resources used for high pressure freezing of worm samples. TEM imaging and analysis were performed in part through the use of the Vanderbilt Cell Imaging Shared Resource (supported by NIH grants CA68485, DK20593, DK58404, DK59637 and EY08126). RNA-sequencing was conducted through the Vanderbilt Technologies for Advanced Genomics VANTAGE core. This work was supported by the Glenn Foundation for Medical Research/American Federation for Aging Research and NIH/NIA R00AG052666 (KB), R01AG073354 (KB).

## Author Contributions

K.B. and G.F. conceived the study; G.F., E.M.R., A.G.M., E.K.F.D., A.H., B.J.C., L.P., and K.B. designed and executed experiments and data analysis; K.B. pursued funding; all authors contributed to manuscript drafting and revision.

## Tables

Supplementary Table 1. Lifespan Statistics

Supplementary Table 2. DE Genes

Supplementary Table 3. Worm Strains

Supplementary Table 4. RNAi

Supplementary Table 5. Reagents and Resources

---

**Extended Data Figure 1:**
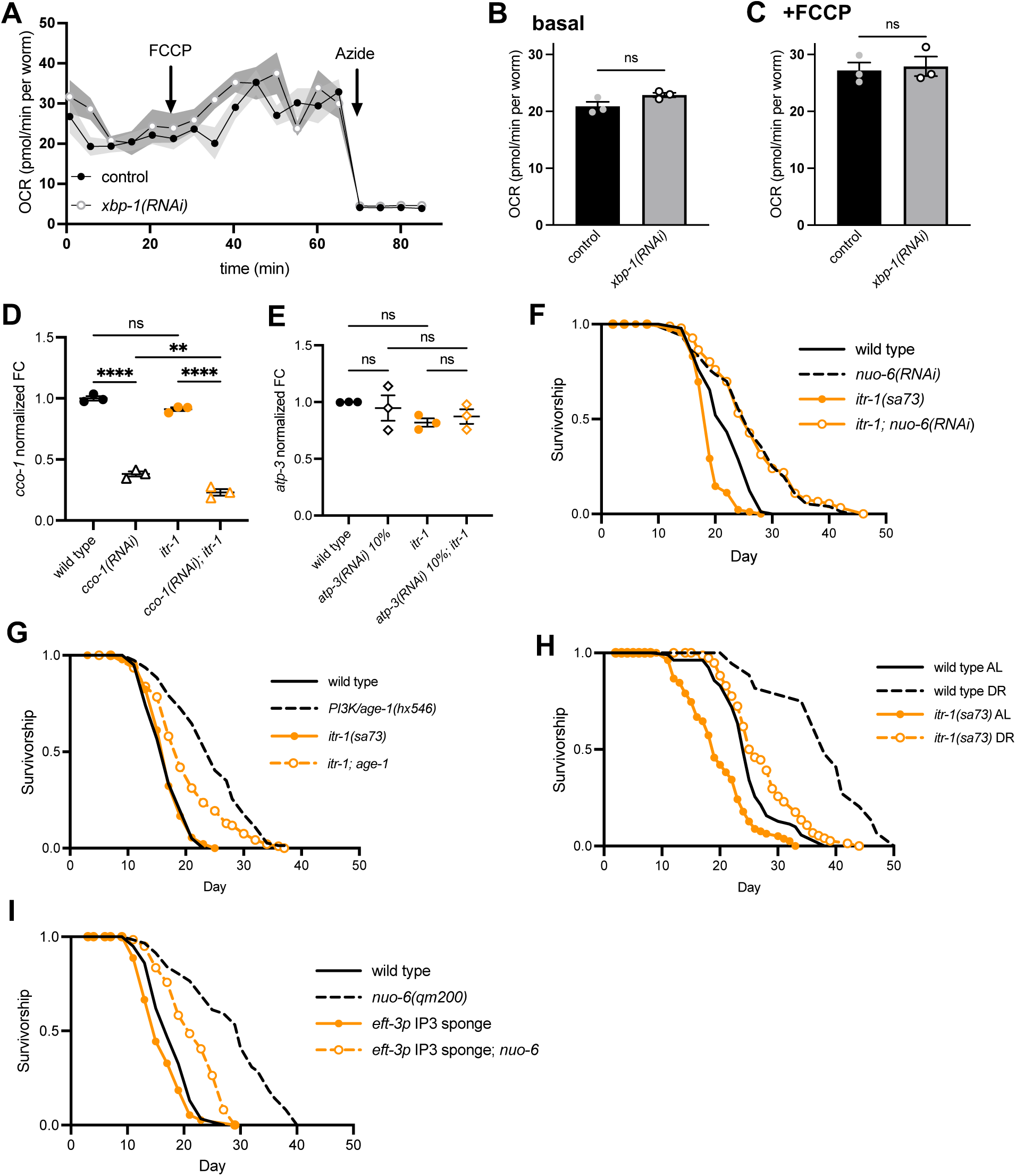
InsP3R requirement for ETC stress adaptation across complexes and longevity contexts. (A) Representative traces of OCR in day 1 wild type adults on control or *xbp-1*(RNAi); mean ± SEM. Additions of FCCP and sodium azide are indicated by arrows. (B, C) Quantification of basal (B) and FCCP-induced (C) OCR; n = 3 independent trials; mean ± SEM; ns p > 0.05 by unpaired t test. (D) Normalized fold change of *cco-1* transcripts from wild type and InsP3R/*itr-1* mutants on *cco-1(RNAi).* n = 3 independent trials; mean ± SEM; one-way ANOVA with Tukey’s multiple comparisons test. (E) Normalized fold change of *atp-3* transcripts from wild type and InsP3R/*itr-1* mutants on 10% *atp-3(RNAi).* n = 3 independent trials; mean ± SEM; one-way ANOVA with Tukey’s multiple comparisons test. (F) Lifespan of InsP3R/*itr-1* mutants on *nuo-6(RNAi)*. n= 100-101 worms per condition. (G) Lifespan of InsP3R/*itr-1* and PI3K/*age-1* mutants. n = 100-102 worms per condition. (H) Lifespan of wild type and InsP3R*/itr-1* worms fed an *ad libitum* or dietary restricted diet. n = 150 worms per condition. (I) Lifespan of ubiquitous *eft-3p* IP3 sponge and Complex I/*nuo-6* double mutants. n = 100-101 worms per condition. Statistical significance of lifespan curves was determined by a log-rank Mantel-Cox test. See Supplementary Table 1 for lifespan statistics. ns p > 0.05; ** p < 0.01; and **** p < 0.0001.

**Extended Data Figure 2:**
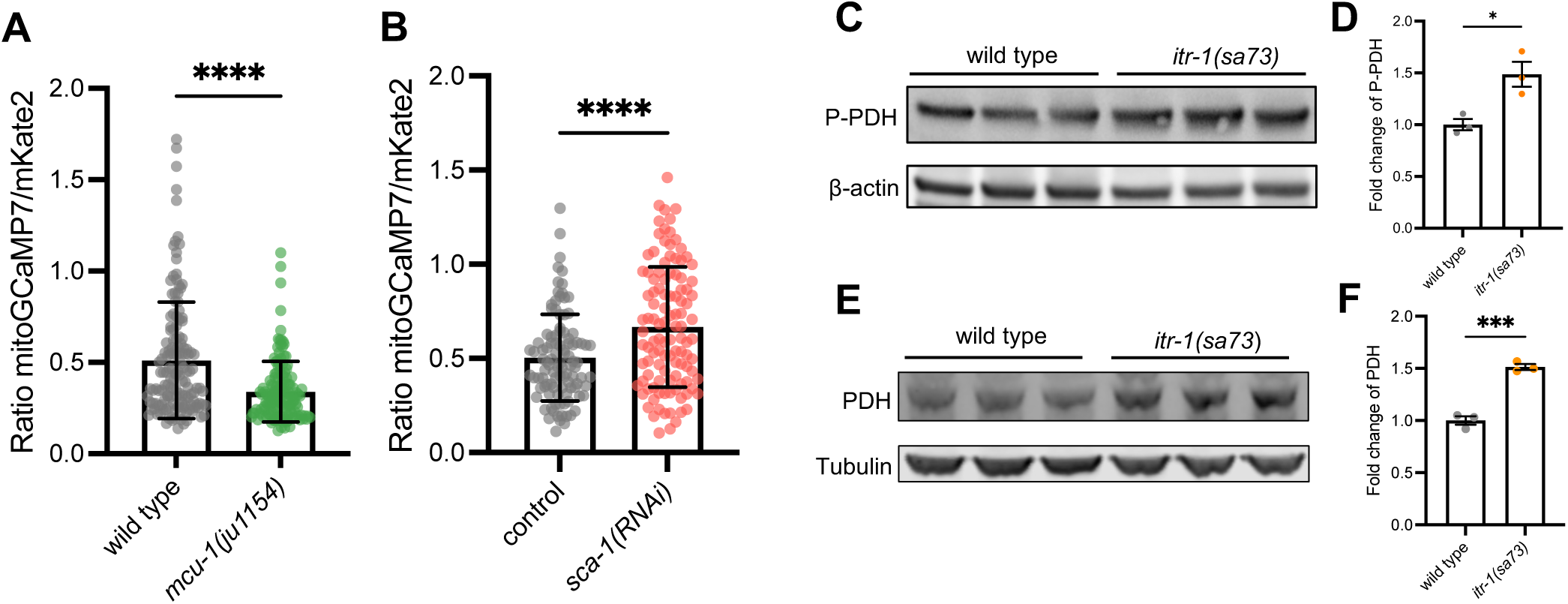
Effects of InsP3R on matrix metabolic processes. (A) Quantification of mitochondrial calcium level indicated by the ratio of GCaMP7:mKate2 fluorescence in intestinal cells of *mcu-1* mutant day 1 adults. n = 146, 140 worms from 4 independent trials (left to right); mean ± SD; **** p < 0.0001 by unpaired t test. (B) Quantification of mitochondrial calcium level indicated by the ratio of GCaMP7:mKate2 fluorescence in intestinal cells of day 1 adults on *sca-1(RNAi)*. n = 102, 111 worms from 3 independent trials (left to right); mean ± SD; **** p < 0.0001 by unpaired t test. (C, D) Western blot (C) and quantification (D) of P-PDH levels in wild type and InsP3R/*itr-1* mutants. n = 3 independent trials; mean ± SEM; p = 0.0212 by unpaired t test. (E, F) Western blot (E) and quantification (F) of total PDH levels in wild type and InsP3R/*itr-1* mutants. n = 3 independent trials; mean ± SEM; p = 0.0005 by unpaired t test.

**Extended Data Figure 3:**
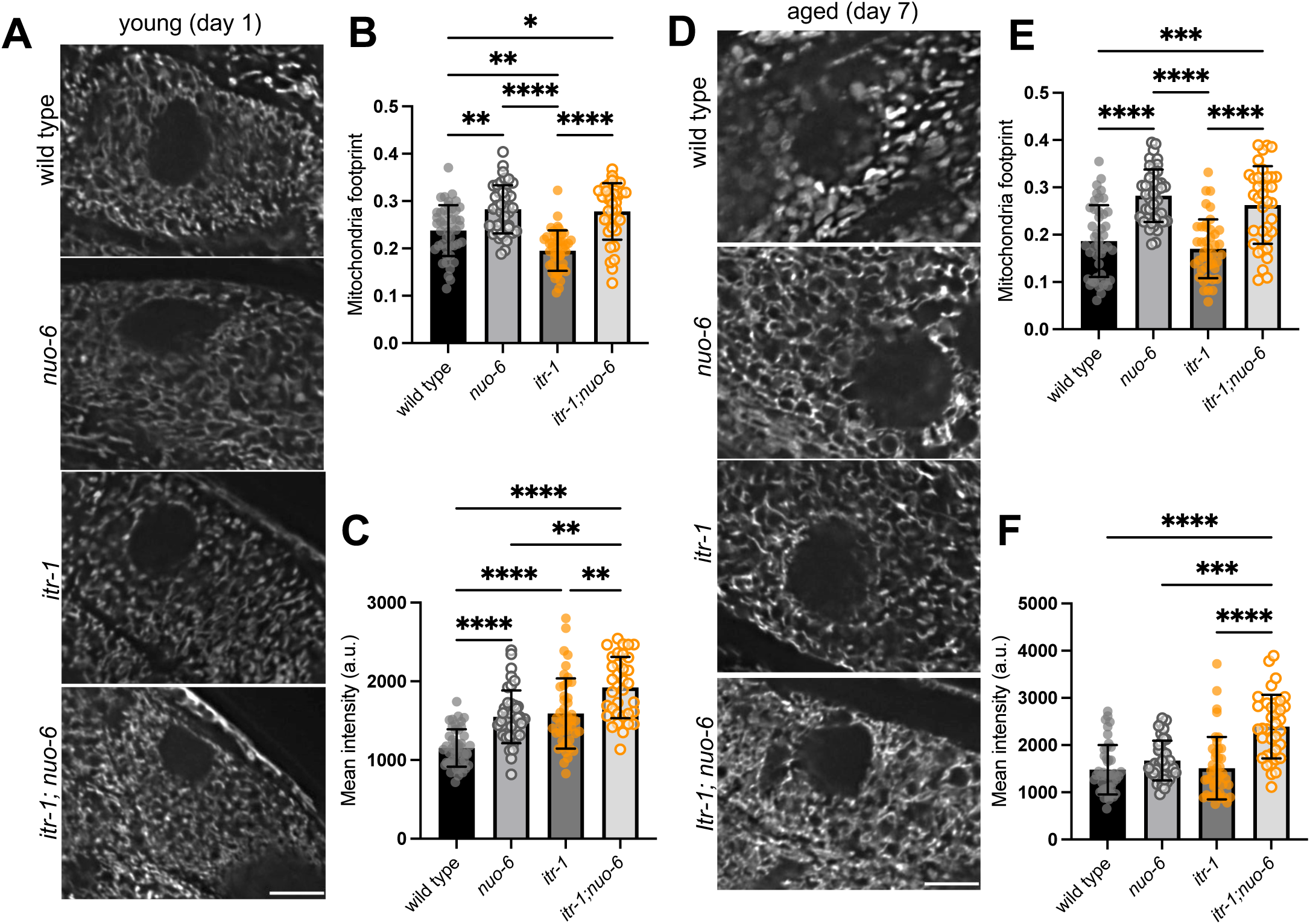
Complex I and InsP3R regulation of intestinal mitochondrial networks. (A) Representative images of mitochondrial networks (COX-4::eGFP) in intestinal cells of young (day 1) worms. Scale bar: 10 µm. (B, C) Quantification of mitochondrial footprint (B) and mean COX-4::eGFP intensity (C) in intestine of young (day 1) worms. n = 46, 41, 44, 35 worms from 2 independent trials (left to right); mean ± SD; Kruskal-Wallis test with Dunn’s multiple comparisons test. (D) Representative images of mitochondrial networks (COX-4::eGFP) in intestinal cells of aged (day 7) worms. Scale bar: 10 µm. (E, F) Quantification of mitochondrial footprint (E) and mean COX-4::eGFP intensity (F) in intestine of aged (day 7) worms. n = 44, 44, 42, 36 worms from 2 independent trials (left to right); mean ± SD; Kruskal-Wallis test with Dunn’s multiple comparisons test. * p < 0.05; ** p < 0.01; *** p < 0.001; and **** p < 0.0001.

**Extended Data Figure 4:**
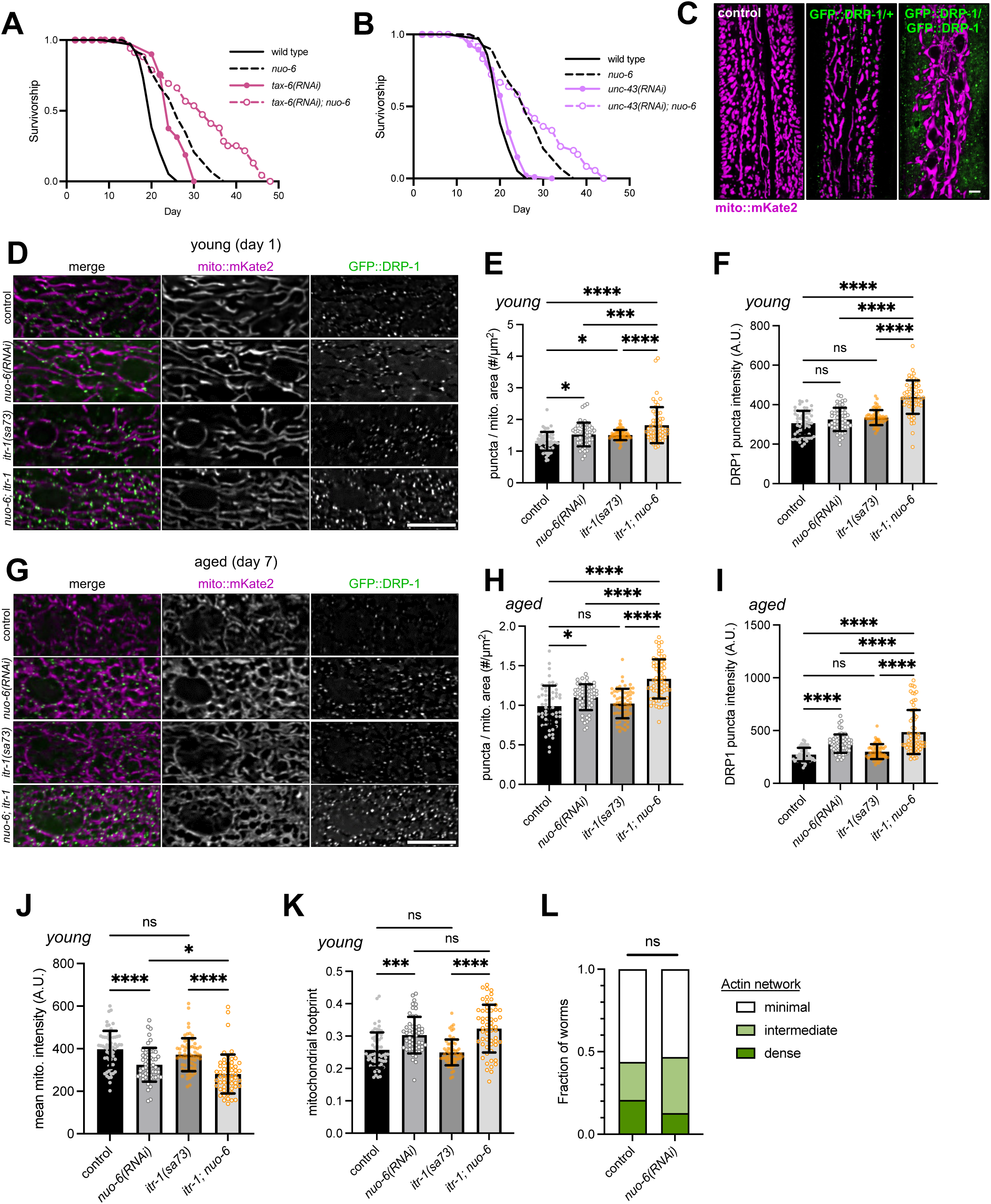
Alterations in fission-associated factors downstream of InsP3R and Complex I. (A) Lifespan of Complex I/*nuo-6* mutants fed *tax-6(RNAi)*. n = 100 worms per condition. (B) Lifespan of Complex I/*nuo-6* mutants fed *unc-43(RNAi)*. n = 100 worms per condition. (C) Representative images of mitochondria (mito::mKate2) and DRP-1 puncta (GFP::DRP-1) in hypodermis of young (day 1) control, GFP::DRP-1 heterozygous, and GFP::DRP-1 homozygous worms. Scale bar: 5 µm. (D) Representative fluorescence images of hypodermal mitochondrial networks (mito::mKate2) and DRP-1 puncta (GFP::DRP-1) in young (day 1) worms. Scale bar: 10 µm. (E) Number of GFP::DRP-1 puncta per mitochondrial area in young (day 1) worms. n = 60, 51, 62, 56 worms from 3 independent trials (left to right); mean ± SD; one-way ANOVA with Tukey’s multiple comparisons test. (F) Mean intensity of GFP::DRP-1 puncta in young (day 1) worms. n = 60, 51, 62, 56 worms from 3 independent trials; mean ± SD; one-way ANOVA with Tukey’s multiple comparisons test. (G) Representative fluorescence images of hypodermal mitochondrial networks (mito::mKate2) and DRP-1 puncta (GFP::DRP-1) in aged (day 7) worms. Scale bar: 10 µm. (H) Number of GFP::DRP-1 puncta per mitochondrial area in aged (day 7) worms. n = 56, 62, 63, 67 worms from 3 independent trials (left to right); mean ± SD; one-way ANOVA with Tukey’s multiple comparisons test. (I) Mean intensity of GFP::DRP-1 puncta in aged (day 7) worms. n = 56, 62, 63, 67 worms from 3 independent trials (left to right); mean ± SD; one-way ANOVA with Tukey’s multiple comparisons test. (J) Quantification of mean mito::mKate2 intensity in hypodermis of young (day 1) worms. n = 60, 51, 62, 56 worms from 3 independent trials; mean ± SD; one-way ANOVA with Tukey’s multiple comparisons test. (K) Quantification of mitochondrial footprint in hypodermis of young (day 1) worms. n = 60, 51, 62, 56 from 2 independent trials (left to right); mean ± SD; one-way ANOVA with Tukey’s multiple comparisons test. (L) Categorization of F-actin network density in worms fed control or *nuo-6(RNAi).* n = 48, 47 worms from 2 independent trials (left to right); p = 0.9177 by Mann-Whitney test. Statistical significance of lifespan curves was determined by a log-rank Mantel-Cox test. See Supplementary Table 1 for lifespan statistics. ns p > 0.05; * p < 0.05; *** p < 0.001; and **** p < 0.0001.

**Extended Data Figure 5:**
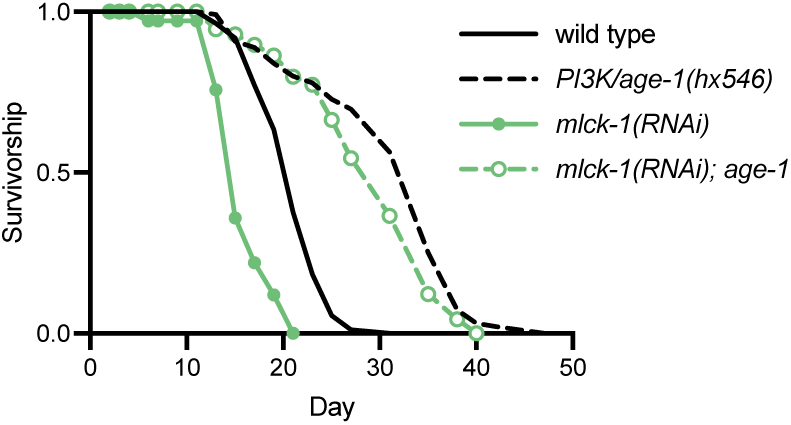
Actomyosin contractility is not required for lifespan extension during reduced insulin signaling. Lifespan of PI3K/*age-1(hx546)* mutants fed *mlck-1(RNAi).* n = 121-223 worms per condition. Statistical significance of lifespan curves was determined by a log-rank Mantel-Cox test. See Supplementary Table 1 for lifespan statistics.

**Extended Data Figure 6:**
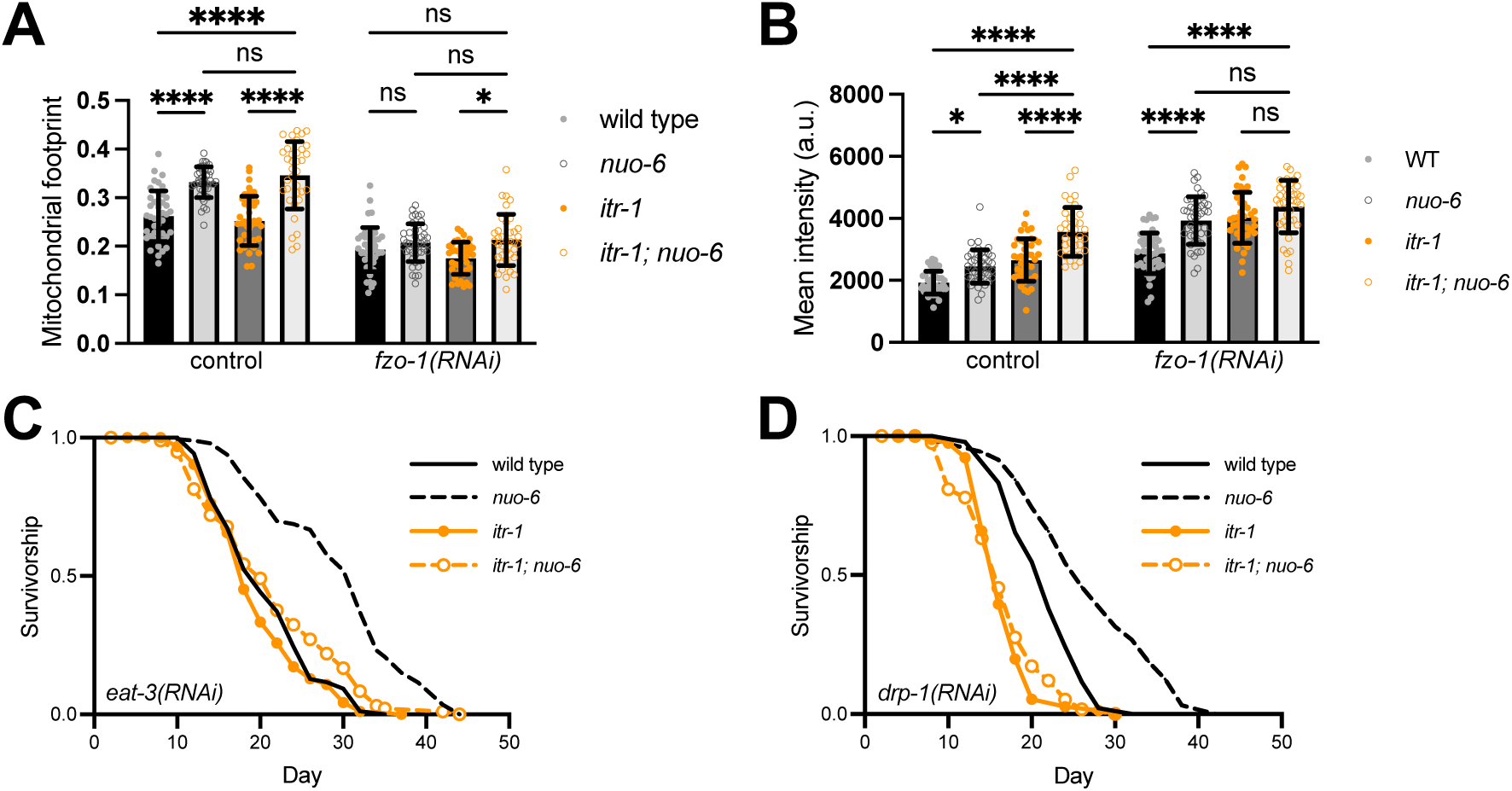
Effects of fission/fusion regulators on lifespan extension. (A, B) Quantification of mitochondrial footprint (A) and mean COX-4::eGFP intensity (B) of hypodermal mitochondrial in young (day 1) worms fed control or *fzo-1(RNAi).* n = 43, 42, 41, 35, 44, 43, 40, 38 worms from 2 independent trials (left to right); mean ± SD; two-way ANOVA with Tukey’s multiple comparisons test. ns p > 0.05; * p < 0.05; **** p < 0.0001. (C, D) Lifespan of InsP3R/*itr-1* and Complex I/*nuo-6* double mutants fed *eat-3(RNAi)* (C) or *drp-1(RNAi)* (D). n = 100 worms per condition. Statistical significance of lifespan curves was determined by a log-rank Mantel-Cox test. See Supplementary Table 1 for lifespan statistics.

## References

1. Ewbank, J.J., Barnes, T.M., Lakowski, B., Lussier, M., Bussey, H., and Hekimi, S. (1997). Structural and functional conservation of the Caenorhabditis elegans timing gene clk-1. Science 275, 980–983.

2. Dillin, A., Hsu, A.-L., Arantes-Oliveira, N., Lehrer-Graiwer, J., Hsin, H., Fraser, A.G., Kamath, R.S., Ahringer, J., and Kenyon, C. (2002). Rates of behavior and aging specified by mitochondrial function during development. Science 298, 2398–2401.

3. Lee, S.S., Lee, R.Y.N., Fraser, A.G., Kamath, R.S., Ahringer, J., and Ruvkun, G. (2003). A systematic RNAi screen identifies a critical role for mitochondria in C. elegans longevity. Nat. Genet. 33, 40–48.

4. Liu, X., Jiang, N., Hughes, B., Bigras, E., Shoubridge, E., and Hekimi, S. (2005). Evolutionary conservation of the clk-1-dependent mechanism of longevity: loss of mclk1 increases cellular fitness and lifespan in mice. Genes Dev. 19, 2424–2434.

5. Dell’agnello, C., Leo, S., Agostino, A., Szabadkai, G., Tiveron, C., Zulian, A., Prelle, A., Roubertoux, P., Rizzuto, R., and Zeviani, M. (2007). Increased longevity and refractoriness to Ca(2+)-dependent neurodegeneration in Surf1 knockout mice. Hum. Mol. Genet. 16, 431–444.

6. Copeland, J.M., Cho, J., Lo, T., Jr, Hur, J.H., Bahadorani, S., Arabyan, T., Rabie, J., Soh, J., and Walker, D.W. (2009). Extension of Drosophila life span by RNAi of the mitochondrial respiratory chain. Curr. Biol. 19, 1591–1598.

7. Barzilai, N., Crandall, J.P., Kritchevsky, S.B., and Espeland, M.A. (2016). Metformin as a Tool to Target Aging. Cell Metab. 23, 1060–1065.

8. Munkácsy, E., and Rea, S.L. (2014). The paradox of mitochondrial dysfunction and extended longevity. Exp. Gerontol. 56, 221–233.

9. Haynes, C.M., and Hekimi, S. (2022). Mitochondrial dysfunction, aging, and the mitochondrial unfolded protein response in Caenorhabditis elegans. Genetics 222. 10.1093/genetics/iyac160.

10. Moehle, E.A., Shen, K., and Dillin, A. (2019). Mitochondrial proteostasis in the context of cellular and organismal health and aging. J. Biol. Chem. 294, 5396–5407.

11. Bennett, C.F., Vander Wende, H., Simko, M., Klum, S., Barfield, S., Choi, H., Pineda, V.V., and Kaeberlein, M. (2014). Activation of the mitochondrial unfolded protein response does not predict longevity in Caenorhabditis elegans. Nat. Commun. 5, 3483.

12. Van Raamsdonk, J.M., Meng, Y., Camp, D., Yang, W., Jia, X., Bénard, C., and Hekimi, S. (2010). Decreased energy metabolism extends life span in Caenorhabditis elegans without reducing oxidative damage. Genetics 185, 559–571.

13. Dues, D.J., Schaar, C.E., Johnson, B.K., Bowman, M.J., Winn, M.E., Senchuk, M.M., and Van Raamsdonk, J.M. (2017). Uncoupling of oxidative stress resistance and lifespan in long-lived isp-1 mitochondrial mutants in Caenorhabditis elegans. Free Radic. Biol. Med. 108, 362–373.

14. Hekimi, S., Lapointe, J., and Wen, Y. (2011). Taking a “good” look at free radicals in the aging process. Trends Cell Biol. 21, 569–576.

15. Rea, S.L. (2005). Metabolism in the Caenorhabditis elegans Mit mutants. Exp. Gerontol. 40, 841–849.

16. Yang, W., and Hekimi, S. (2010). Two modes of mitochondrial dysfunction lead independently to lifespan extension in Caenorhabditis elegans. Aging Cell 9, 433–447.

17. Loubiere, C., Clavel, S., Gilleron, J., Harisseh, R., Fauconnier, J., Ben-Sahra, I., Kaminski, L., Laurent, K., Herkenne, S., Lacas-Gervais, S., et al. (2017). The energy disruptor metformin targets mitochondrial integrity via modification of calcium flux in cancer cells. Sci. Rep. 7, 5040.

18. Balderas, E., Eberhardt, D.R., Lee, S., Pleinis, J.M., Sommakia, S., Balynas, A.M., Yin, X., Parker, M.C., Maguire, C.T., Cho, S., et al. (2022). Mitochondrial calcium uniporter stabilization preserves energetic homeostasis during Complex I impairment. Nat. Commun. 13, 2769.

19. Jaña, F., Bustos, G., Rivas, J., Cruz, P., Urra, F., Basualto-Alarcón, C., Sagredo, E., Ríos, M., Lovy, A., Dong, Z., et al. (2019). Complex I and II are required for normal mitochondrial Ca2+ homeostasis. Mitochondrion 49, 73–82.

20. Rossi, A., Pizzo, P., and Filadi, R. (2019). Calcium, mitochondria and cell metabolism: A functional triangle in bioenergetics. Biochim. Biophys. Acta Mol. Cell Res. 1866, 1068–1078.

21. Hajnóczky, G., Hager, R., and Thomas, A.P. (1999). Mitochondria suppress local feedback activation of inositol 1,4, 5-trisphosphate receptors by Ca2+. J. Biol. Chem. 274, 14157–14162.

22. Alzheimer’s Association Calcium Hypothesis Workgroup (2017). Calcium Hypothesis of Alzheimer’s disease and brain aging: A framework for integrating new evidence into a comprehensive theory of pathogenesis. Alzheimers. Dement. 13, 178-182.e17.

23. Mattson, M.P., and Arumugam, T.V. (2018). Hallmarks of Brain Aging: Adaptive and Pathological Modification by Metabolic States. Cell Metab. 27, 1176–1199.

24. Csordás, G., Thomas, A.P., and Hajnóczky, G. (1999). Quasi-synaptic calcium signal transmission between endoplasmic reticulum and mitochondria. EMBO J. 18, 96–108.

25. Cárdenas, C., Miller, R.A., Smith, I., Bui, T., Molgó, J., Müller, M., Vais, H., Cheung, K.-H., Yang, J., Parker, I., et al. (2010). Essential regulation of cell bioenergetics by constitutive InsP3 receptor Ca2+ transfer to mitochondria. Cell 142, 270–283.

26. Lovy, A., Foskett, J.K., and Cárdenas, C. (2016). InsP3R, the calcium whisperer: Maintaining mitochondrial function in cancer. Mol Cell Oncol 3, e1185563.

27. Denton, R.M. (2009). Regulation of mitochondrial dehydrogenases by calcium ions. Biochim. Biophys. Acta 1787, 1309–1316.

28. Sharma, A., Smith, H.J., Yao, P., and Mair, W.B. (2019). Causal roles of mitochondrial dynamics in longevity and healthy aging. EMBO Rep. 20, e48395.

29. Cereghetti, G.M., Stangherlin, A., Martins de Brito, O., Chang, C.R., Blackstone, C., Bernardi, P., and Scorrano, L. (2008). Dephosphorylation by calcineurin regulates translocation of Drp1 to mitochondria. Proc. Natl. Acad. Sci. U. S. A. 105, 15803–15808.

30. Chakrabarti, R., Ji, W.-K., Stan, R.V., de Juan Sanz, J., Ryan, T.A., and Higgs, H.N. (2018). INF2-mediated actin polymerization at the ER stimulates mitochondrial calcium uptake, inner membrane constriction, and division. J. Cell Biol. 217, 251–268.

31. Fung, T.S., Chakrabarti, R., Kollasser, J., Rottner, K., Stradal, T.E.B., Kage, F., and Higgs, H.N. (2022). Parallel kinase pathways stimulate actin polymerization at depolarized mitochondria. Curr. Biol. 32, 1577–1592.e8.

32. Kruppa, A.J., Kishi-Itakura, C., Masters, T.A., Rorbach, J.E., Grice, G.L., Kendrick-Jones, J., Nathan, J.A., Minczuk, M., and Buss, F. (2018). Myosin VI-Dependent Actin Cages Encapsulate Parkin-Positive Damaged Mitochondria. Dev. Cell 44, 484–499.e6.

33. Hsieh, C.-W., and Yang, W.Y. (2019). Omegasome-proximal PtdIns(4,5)P2 couples F-actin mediated mitoaggregate disassembly with autophagosome formation during mitophagy. Nat. Commun. 10, 969.

34. Chakrabarti, R., Fung, T.S., Kang, T., Elonkirjo, P.W., Suomalainen, A., Usherwood, E.J., and Higgs, H.N. (2022). Mitochondrial dysfunction triggers actin polymerization necessary for rapid glycolytic activation. J. Cell Biol. 221. 10.1083/jcb.202201160.

35. Iwasa, H., Yu, S., Xue, J., and Driscoll, M. (2010). Novel EGF pathway regulators modulate C. elegans healthspan and lifespan via EGF receptor, PLC-gamma, and IP3R activation. Aging Cell 9, 490–505.

36. Burkewitz, K., Feng, G., Dutta, S., Kelley, C.A., Steinbaugh, M., Cram, E.J., and Mair, W.B. (2020). Atf-6 Regulates Lifespan through ER-Mitochondrial Calcium Homeostasis. Cell Rep. 32, 108125.

37. Walker, D.S., Gower, N.J.D., Ly, S., Bradley, G.L., and Baylis, H.A. (2002). Regulated disruption of inositol 1,4,5-trisphosphate signaling in Caenorhabditis elegans reveals new functions in feeding and embryogenesis. Mol. Biol. Cell 13, 1329–1337.

38. Espelt, M.V., Estevez, A.Y., Yin, X., and Strange, K. (2005). Oscillatory Ca2+ signaling in the isolated Caenorhabditis elegans intestine: role of the inositol-1,4,5-trisphosphate receptor and phospholipases C beta and gamma. J. Gen. Physiol. 126, 379–392.

39. Kovacevic, I., Orozco, J.M., and Cram, E.J. (2013). Filamin and phospholipase C-ε are required for calcium signaling in the Caenorhabditis elegans spermatheca. PLoS Genet. 9, e1003510.

40. Cardenas, C., Lovy, A., Silva-Pavez, E., Urra, F., Mizzoni, C., Ahumada-Castro, U., Bustos, G., Jaňa, F., Cruz, P., Farias, P., et al. (2020). Cancer cells with defective oxidative phosphorylation require endoplasmic reticulum-to-mitochondria Ca2+ transfer for survival. Sci. Signal. 13. 10.1126/scisignal.aay1212.

41. Young, M.P., Schug, Z.T., Booth, D.M., Yule, D.I., Mikoshiba, K., Hajnόczky, G., and Joseph, S.K. (2022). Metabolic adaptation to the chronic loss of Ca2+ signaling induced by KO of IP3 receptors or the mitochondrial Ca2+ uniporter. J. Biol. Chem. 298, 101436.

42. Rea, S.L., Ventura, N., and Johnson, T.E. (2007). Relationship between mitochondrial electron transport chain dysfunction, development, and life extension in Caenorhabditis elegans. PLoS Biol. 5, e259.

43. Hajnóczky, G., Robb-Gaspers, L.D., Seitz, M.B., and Thomas, A.P. (1995). Decoding of cytosolic calcium oscillations in the mitochondria. Cell 82, 415–424.

44. Cho, B., Cho, H.M., Jo, Y., Kim, H.D., Song, M., Moon, C., Kim, H., Kim, K., Sesaki, H., Rhyu, I.J., et al. (2017). Constriction of the mitochondrial inner compartment is a priming event for mitochondrial division. Nat. Commun. 8, 15754.

45. Holdorf, A.D., Higgins, D.P., Hart, A.C., Boag, P.R., Pazour, G.J., Walhout, A.J.M., and Walker, A.K. (2020). WormCat: An Online Tool for Annotation and Visualization of Caenorhabditis elegans Genome-Scale Data. Genetics 214, 279–294.

46. Nargund, A.M., Pellegrino, M.W., Fiorese, C.J., Baker, B.M., and Haynes, C.M. (2012). Mitochondrial import efficiency of ATFS-1 regulates mitochondrial UPR activation. Science 337, 587–590.

47. Nargund, A.M., Fiorese, C.J., Pellegrino, M.W., Deng, P., and Haynes, C.M. (2015). Mitochondrial and nuclear accumulation of the transcription factor ATFS-1 promotes OXPHOS recovery during the UPR(mt). Mol. Cell 58, 123–133.

48. Wu, Z., Senchuk, M.M., Dues, D.J., Johnson, B.K., Cooper, J.F., Lew, L., Machiela, E., Schaar, C.E., DeJonge, H., Blackwell, T.K., et al. (2018). Mitochondrial unfolded protein response transcription factor ATFS-1 promotes longevity in a long-lived mitochondrial mutant through activation of stress response pathways. BMC Biol. 16, 147.

49. Liu, Y., Samuel, B.S., Breen, P.C., and Ruvkun, G. (2014). Caenorhabditis elegans pathways that surveil and defend mitochondria. Nature 508, 406–410.

50. Haeussler, S., Yeroslaviz, A., Rolland, S.G., Luehr, S., Lambie, E.J., and Conradt, B. (2021). Genome-wide RNAi screen for regulators of UPRmt in Caenorhabditis elegans mutants with defects in mitochondrial fusion. G3 11. 10.1093/g3journal/jkab095.

51. Xu, S., Wang, P., Zhang, H., Gong, G., Gutierrez Cortes, N., Zhu, W., Yoon, Y., Tian, R., and Wang, W. (2016). CaMKII induces permeability transition through Drp1 phosphorylation during chronic β-AR stimulation. Nat. Commun. 7, 13189.

52. Dong, M.-Q., Venable, J.D., Au, N., Xu, T., Park, S.K., Cociorva, D., Johnson, J.R., Dillin, A., and Yates, J.R., 3rd (2007). Quantitative mass spectrometry identifies insulin signaling targets in C. elegans. Science 317, 660–663.

53. Tao, L., Xie, Q., Ding, Y.-H., Li, S.-T., Peng, S., Zhang, Y.-P., Tan, D., Yuan, Z., and Dong, M.-Q. (2013). CAMKII and calcineurin regulate the lifespan of Caenorhabditis elegans through the FOXO transcription factor DAF-16. Elife 2, e00518.

54. Montecinos-Franjola, F., Bauer, B.L., Mears, J.A., and Ramachandran, R. (2020). GFP fluorescence tagging alters dynamin-related protein 1 oligomerization dynamics and creates disassembly-refractory puncta to mediate mitochondrial fission. Sci. Rep. 10, 14777.

55. Fung, T.S., Chakrabarti, R., and Higgs, H.N. (2023). The multiple links between actin and mitochondria. Nat. Rev. Mol. Cell Biol. 24, 651–667.

56. Heissler, S.M., and Sellers, J.R. (2014). Myosin light chains: Teaching old dogs new tricks. Bioarchitecture 4, 169–188.

57. Korobova, F., Ramabhadran, V., and Higgs, H.N. (2013). An actin-dependent step in mitochondrial fission mediated by the ER-associated formin INF2. Science 339, 464–467.

58. Rambold, A.S., Kostelecky, B., Elia, N., and Lippincott-Schwartz, J. (2011). Tubular network formation protects mitochondria from autophagosomal degradation during nutrient starvation. Proc. Natl. Acad. Sci. U. S. A. 108, 10190–10195.

59. Twig, G., Elorza, A., Molina, A.J.A., Mohamed, H., Wikstrom, J.D., Walzer, G., Stiles, L., Haigh, S.E., Katz, S., Las, G., et al. (2008). Fission and selective fusion govern mitochondrial segregation and elimination by autophagy. EMBO J. 27, 433–446.

60. Pandya, J.D., Grondin, R., Yonutas, H.M., Haghnazar, H., Gash, D.M., Zhang, Z., and Sullivan, P.G. (2015). Decreased mitochondrial bioenergetics and calcium buffering capacity in the basal ganglia correlates with motor deficits in a nonhuman primate model of aging. Neurobiol. Aging 36, 1903–1913.

61. Filadi, R., Leal, N.S., Schreiner, B., Rossi, A., Dentoni, G., Pinho, C.M., Wiehager, B., Cieri, D., Calì, T., Pizzo, P., et al. (2018). TOM70 Sustains Cell Bioenergetics by Promoting IP3R3-Mediated ER to Mitochondria Ca2+ Transfer. Curr. Biol. 28, 369–382.e6.

62. Mallilankaraman, K., Cárdenas, C., Doonan, P.J., Chandramoorthy, H.C., Irrinki, K.M., Golenár, T., Csordás, G., Madireddi, P., Yang, J., Müller, M., et al. (2012). MCUR1 is an essential component of mitochondrial Ca2+ uptake that regulates cellular metabolism. Nat. Cell Biol. 14, 1336–1343.

63. Pan, X., Liu, J., Nguyen, T., Liu, C., Sun, J., Teng, Y., Fergusson, M.M., Rovira, I.I., Allen, M., Springer, D.A., et al. (2013). The physiological role of mitochondrial calcium revealed by mice lacking the mitochondrial calcium uniporter. Nat. Cell Biol. 15, 1464–1472.

64. Kosmach, A., Roman, B., Sun, J., Femnou, A., Zhang, F., Liu, C., Combs, C.A., Balaban, R.S., and Murphy, E. (2021). Monitoring mitochondrial calcium and metabolism in the beating MCU-KO heart. Cell Rep. 37, 109846.

65. Szibor, M., Gizatullina, Z., Gainutdinov, T., Endres, T., Debska-Vielhaber, G., Kunz, M., Karavasili, N., Hallmann, K., Schreiber, F., Bamberger, A., et al. (2020). Cytosolic, but not matrix, calcium is essential for adjustment of mitochondrial pyruvate supply. J. Biol. Chem. 295, 4383–4397.

66. Glancy, B., and Balaban, R.S. (2012). Role of mitochondrial Ca2+ in the regulation of cellular energetics. Biochemistry 51, 2959–2973.

67. Pérez-Liébana, I., Juaristi, I., González-Sánchez, P., González-Moreno, L., Rial, E., Podunavac, M., Zakarian, A., Molgó, J., Vallejo-Illarramendi, A., Mosqueira-Martín, L., et al. (2022). A Ca2+-Dependent Mechanism Boosting Glycolysis and OXPHOS by Activating Aralar-Malate-Aspartate Shuttle, upon Neuronal Stimulation. J. Neurosci. 42, 3879–3895.

68. Roy Chowdhury, A., Srinivasan, S., Csordás, G., Hajnóczky, G., and Avadhani, N.G. (2020). Dysregulation of RyR Calcium Channel Causes the Onset of Mitochondrial Retrograde Signaling. iScience 23, 101370.

69. Meisel, J.D., Miranda, M., Skinner, O.S., Wiesenthal, P.P., Wellner, S.M., Jourdain, A.A., Ruvkun, G., and Mootha, V.K. (2024). Hypoxia and intra-complex genetic suppressors rescue complex I mutants by a shared mechanism. Cell 187, 659–675.e18.

70. Kim, E., Sun, L., Gabel, C.V., and Fang-Yen, C. (2013). Long-term imaging of Caenorhabditis elegans using nanoparticle-mediated immobilization. PLoS One 8, e53419.

71. Koopman, M., Michels, H., Dancy, B.M., Kamble, R., Mouchiroud, L., Auwerx, J., Nollen, E.A.A., and Houtkooper, R.H. (2016). A screening-based platform for the assessment of cellular respiration in Caenorhabditis elegans. Nat. Protoc. 11, 1798–1816.

72. Ching, T.-T., and Hsu, A.-L. (2011). Solid plate-based dietary restriction in Caenorhabditis elegans. J. Vis. Exp. 10.3791/2701.

73. Dobin, A., Davis, C.A., Schlesinger, F., Drenkow, J., Zaleski, C., Jha, S., Batut, P., Chaisson, M., and Gingeras, T.R. (2013). STAR: ultrafast universal RNA-seq aligner. Bioinformatics 29, 15–21.

74. Bélanger, S., Berensmann, H., Baena, V., Duncan, K., Meyers, B.C., Narayan, K., and Czymmek, K.J. (2022). A versatile enhanced freeze-substitution protocol for volume electron microscopy. Front Cell Dev Biol 10, 933376.

